# Geometry-Constrained Prediction of Catalytically Competent Kinase Domains Across the Human Kinome

**DOI:** 10.1101/2025.11.03.686188

**Authors:** Yuxuan Wang, Qi Hu, Jing Huang

## Abstract

Understanding cellular signaling networks benefits from accurate structural modeling of protein kinases across their different functional states. However, most human kinases lack experimentally determined active-state structures, and existing computational methods often fail to reliably predict catalytically competent conformations. Here, we present a geometry-constrained modeling framework based on AFEXplorer (AFEX) that enables robust prediction of active, ATP- and magnesium-bound states for 436 catalytic kinase domains across the human kinome. By explicitly enforcing key biophysical criteria, AFEX overcomes the template dependence and conformational bias inherent in AlphaFold2 and AlphaFold3. Benchmarking against 156 kinase-substrate complexes shows that AFEX achieves sub-2□Å backbone RMSD in 99% of cases and accurately recapitulates critical catalytic features with precise ATP coordination. Notably, AFEX correctly predicted active conformations for EGFR and CDK2, where AlphaFold3 failed. These improvements are consistent across diverse kinase families, including both well-characterized kinases and those underrepresented or absent from AlphaFold’s training data. AFEX bridges a critical gap in kinase biology by providing a generalizable framework for studying active conformations and dynamics in kinases and other signaling proteins.

## Introduction

Protein kinases are central regulators of intracellular phosphorylation signaling, orchestrating over 30% of cellular protein activity and governing critical processes such as proliferation, differentiation, and apoptosis ^1^. The human kinome comprises 481 genes, each encoding at least one canonical full-length kinase domain; 13 of these genes contain two kinase domains, resulting in a total of 494 kinase domains ^2,3^. Of these, 437 are predicted to be catalytically active and mediate the phosphorylation of serine, threonine, or tyrosine residues to modulate signaling networks ^4^. These kinases are classified into eight major families (AGC, CAMK, CK1, CMGC, STE, TK, TKL, and RGC), each comprising distinct subfamilies with specialized roles in processes ranging from transcriptional regulation to cell cycle control ^3^. All kinases share a conserved catalytic core, which includes an N-terminal β-strand lobe with a critical αC-helix and a C-terminal α-helical lobe ^5^. Crucially, catalytic activity depends on precise and uniform geometric arrangements: the activation loop must adopt an extended conformation, and the αC-helix must rotate inward to align the ATP-binding and substrate recognition sites ^6,7^. In contrast, inactive states exhibit structural heterogeneity, such as DFG motif rearrangements or αC-helix displacement, which alter the accessibility of the binding pocket and impair catalytic function ^7^.

Structural characterization of active-state conformations across the catalytic human kinome is crucial for elucidating the biochemical mechanisms underlying kinase signaling. Conformational dynamics regulate catalytic activity, substrate recognition, allosteric control, and subcellular localization, collectively coordinating cellular responses to extracellular stimuli. In general, kinases act as molecular switches, cycling between active and inactive states through conformational changes in key structural elements such as the activation segment, αC-helix, and DFG motif. Disruptions to these regions, induced by mutations or inhibitors, can lead to aberrant signaling and are implicated in diseases such as cancer, inflammation, and neurodegeneration ^8^. For example, NMR studies of Abelson kinase (Abl) demonstrate that the active conformation predominates (∼88%) under physiological conditions, and drug-resistant mutations shift the conformational equilibrium to further stabilize the active state ^5^.

The transition into the active-state conformation is sometimes coupled to the relocation of kinases and changes in their binding partners, yet the atomistic basis and mechanistic understanding of these processes remain poorly defined. For example, activation of Protein Kinase D (PKD) is essential for the fission of transport carriers from the trans-Golgi network (TGN) to the plasma membrane. Kinase-inactive PKD-K618N mutants accumulate at the TGN, resulting in extensive membrane tubulation and defective vesicle release ^9,10^. Activation of ERK1 triggers its nuclear translocation, marking the final step of the MAPK signaling cascade, where it engages transcription factors to regulate cell cycle progression ^11^. Such activation-induced subcellular relocalization is a conserved regulatory mechanism: during apoptosis, PKCδ translocates to the nucleus to promote lamin B1 disassembly ^12^, whereas PKCε enhances ERK nuclear import ^13^, highlighting the functional significance of active-state conformations. Accurately capturing these active-state structures can therefore help elucidate the spatial and functional regulation of kinase signaling.

Despite the conservation of core activation motifs across the kinome, structural diversity precludes straightforward generalization. A major challenge in characterizing active-state conformations lies in the limited availability of experimentally resolved active kinase structures. Among 272 unique kinase domains deposited in the Protein Data Bank (PDB), only 156 meet established criteria for active states, and many lack complete resolution of the activation loop ^14^. This paucity stems in part from inherent limitations of traditional structural biology techniques, such as X-ray crystallography and cryo-electron microscopy, which face challenges in sample preparation and data collection when resolving catalytically active conformations, particularly for low molecular weight kinases ^15^. Deep learning–based methods, notably AlphaFold, have transformed protein structure prediction ^16– 18^, yet due to biases in training data, these models often fail to capture functionally relevant conformational states, including the active conformations of kinases. For instance, only 48% of catalytic domains in the EBI dataset exhibit active-like geometry ^18^. Although AlphaFold-predicted structures have shown promise in facilitating ligand discovery ^19^, their utility is constrained by structural uncertainties in low-confidence regions, inaccuracies in side-chain modeling, and substantial variations in binding pocket volumes ^20–22^, underscoring the need for accurate, functionally relevant structural models.

Methods that integrate physics-based enhanced sampling techniques to generate diverse conformations have been proposed to improve the efficiency and selectivity of structure-guided drug design, but these require computationally expensive molecular dynamics (MD) simulations and rigorous validation ^23^. Existing template-based approaches for predicting kinome-wide active-like kinase conformations, although informative, lack benchmarkability because predictions were generated using exclusively active-state templates from the PDB—structures that typically serve as ground truth in evaluation frameworks ^4,14^. A more widely adopted strategy for extracting alternative conformations from AlphaFold relies on subsampling multiple sequence alignment (MSA) inputs. Reducing MSA depth has been shown to enhance structural diversity and accuracy in AlphaFold2 models, illustrating the algorithm’s capacity to sample multiple conformational states of a given protein ^24^. Several heuristic methods have since emerged to improve conformational sampling for alternative conformation prediction (ACP). The AF-cluster approach ^25^ exploits clustering of MSA sequences by similarity to enable AlphaFold2 to capture alternative states of metamorphic proteins. AFsample2 ^26^ introduces random masking of MSA columns to attenuate co-evolutionary signals, thereby increasing structural diversity in AlphaFold2-generated ensembles and enabling effective ACP. Guan et al. proposed using the concept of frustration from the energy landscape theory to guide MSA subsampling for predicting conformational structures and motions ^27^. Xing et al. developed AF-ClaSeq, which isolates specific co-evolutionary signals through sequence purification by iteratively generating MSA subsets, performing structure predictions, and comparing them to previously sampled conformations to identify novel states ^28^. Concurrently, we developed a framework, AFEXplorer (AFEX), that integrates collective variables (CVs)—a core concept in enhanced sampling methodologies—into the challenge of ACP. By reformulating ACP as an optimization problem, AFEX employs a CV-defined loss function to guide conditioned conformational sampling ^29^.

In this work, we extend this geometry-driven framework to harness AlphaFold2 for predicting active-state conformations of kinases. By integrating critical structural constraints with refined MSA processing, AFEX successfully sampled active-state models for 436 catalytic kinase domains, with MASTL excluded due to its long activation loop. Benchmarking against recently resolved experimental structures reveals that AFEX achieves substantial improvement over both AlphaFold2 and AlphaFold3 in accurately capturing these functionally crucial conformations. This systematic prediction of kinase activation states opens new avenues for mechanistic insights and has the potential to accelerate structure-based drug discovery across the human kinome.

## Results

### AFEX Enables Prediction of Active-State Conformations and Ligand Imputation for 436 Human Kinase Domains

Active-state conformations were successfully sampled for 436 catalytic domains across 429 human protein kinases using AFEX optimization, the fundamental geometric constraints guiding the sampling were derived from the structural analysis of all kinase–substrate complexes available in the PDB ^4^. The general active-like conformation was defined by the following constraints: a DFG-in conformation, a BLAminus conformation of the XDFG motif, preservation of the salt bridge between the C-helix and β3 strand, and proper extension of the activation loop with contact to the catalytic loop (see Methods). These criteria were validated through our statistical analysis of the same geometric parameters across all ATP- and ADP-bound kinase structures in the PDB (**Supplementary Figure 1**).

During the AFEX optimization process, a collective loss function was incorporated into the AlphaFold2 inference module, integrating geometric constraints with confidence metrics (pLDDT) for both the global structure and the activation segment (**Figure 1**). Gradient descent was utilized to refine the MSA representation features within AlphaFold2, driving convergence toward conformations that minimized the CV-defined loss. Subsequent statistical analysis of AFEX-predicted structures revealed that both global and activation segment-specific pLDDT scores exceeded 0.95, consistently meeting criteria indicative of the active state (**Supplementary Figure 2**).

**Figure 1.**
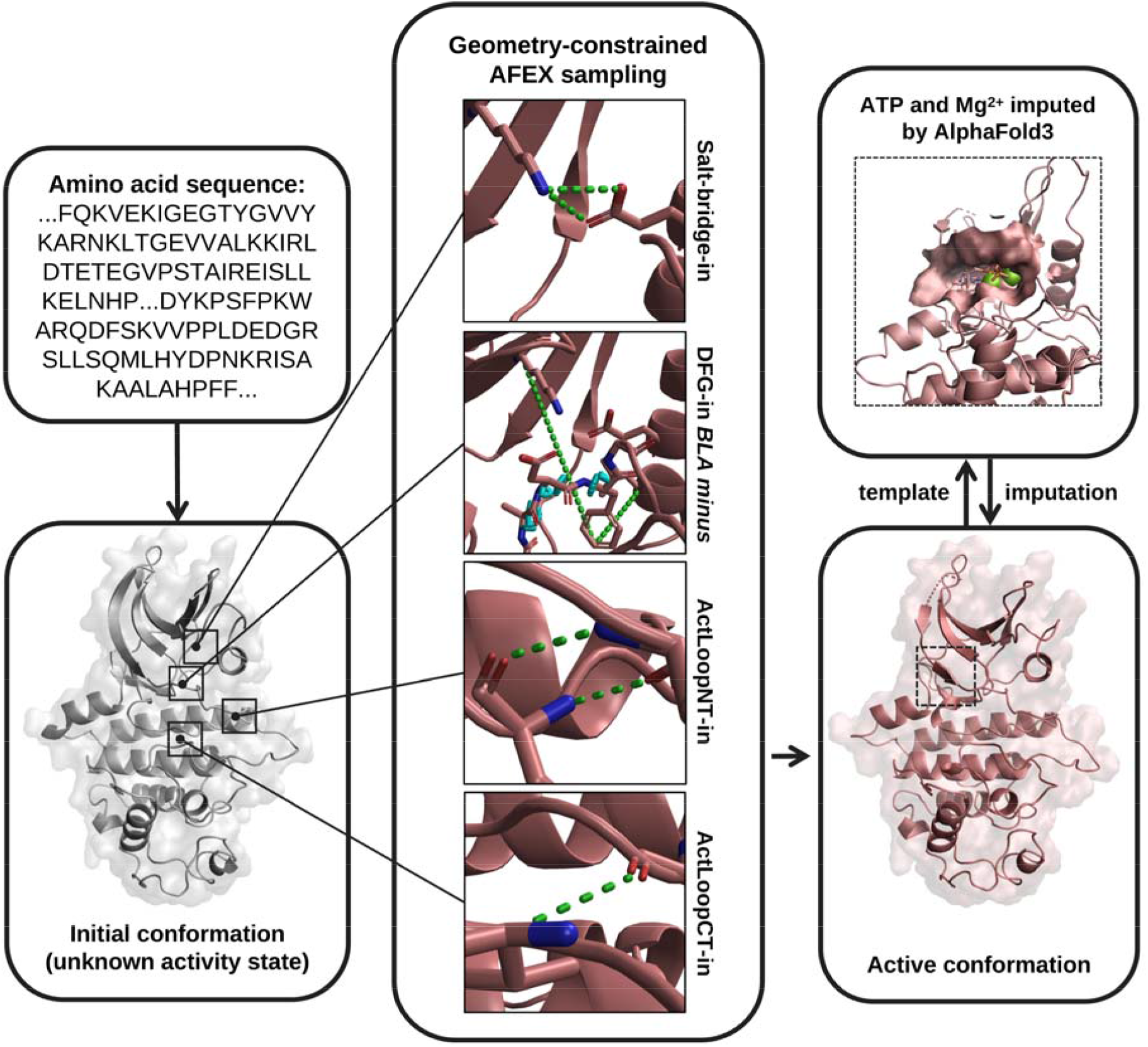
Schematic overview of the AFEXplorer (AFEX) workflow for geometry-constrained prediction of active-state conformations of 436 human kinase domains across the kinome. AFEX systematically samples the conformational space of 436 human kinase domains by iteratively refining MSA representations generated by AlphaFold2. The refinement incorporates explicit geometric constraints that enforce a DFG-in conformation, a BLA-minus conformation of the XDFG motif, preservation of the salt bridge between the C-helix and the β3 strand, and proper extension of the activation loop with contact to the catalytic loop (see Methods). Green dashed lines denote distance constraints imposed between key residues, while cyan dashed lines indicate dihedral angle constraints used to guide critical backbone torsions toward catalytically competent geometries.

We then used the AFEX-predicted active-state structure for each kinase as the sole template in an AlphaFold3 prediction to impute ATP and magnesium ions. The structures were aligned before and after ligand incorporation, and the coordinates of ATP and the two Mg^2^□ ions were transferred from the AlphaFold3 output to the corresponding AFEX structure (see Methods). Minor atomic clashes were resolved through energy minimization in Chimera, yielding AFEX-imputed structures with RMSD values below 0.25 Å relative to the original AFEX predictions (**Supplementary Figure 2e**).

### AFEX Outperforms AlphaFold2 and AlphaFold3 in Predicting Active and ATP-Bound Kinase Conformations

To quantify model performance, we define accuracy as the proportion of kinase models whose backbone root mean square deviation (RMSD) falls below a specified threshold relative to experimentally determined reference structures. We report accuracy at two thresholds, RMSD < 1 Å and RMSD < 2 Å, and evaluate it separately for the entire catalytic domain and for the activation loop (defined as the segment spanning the DFG to APE motifs), which is a highly flexible region critical for kinase function. This allows us to distinguish overall structural agreement from accuracy in the most functionally relevant conformational element.

We benchmarked AFEX models against a curated dataset of 156 experimentally determined substrate-bound active-state kinase structures, selected from 3,010 PDB entries, to evaluate their predictive accuracy. The ground truth dataset was stratified by release date relative to the AlphaFold2 and AlphaFold3 training cutoffs. Systematic comparisons were conducted between models generated by AFEX and those predicted by AlphaFold2 and AlphaFold3 to ensure a rigorous and unbiased statistical assessment.

For comparison with AlphaFold2 models trained on PDB data available prior to April 2019, accuracy was assessed on a subset of 145 substrate-bound active-state kinases. AFEX achieved an average RMSD <1 Å for 66.9% of activation segments and 75.9% for global catalytic domains (**Figure 2a and Supplementary Figure 3a**). With the exception of CSF1R, all kinases predicted by AFEX demonstrated RMSDs <2 Å in both their global catalytic domains and activation segments, exhibited higher prediction accuracy for activation segments than Alphafold2 (99.3% vs. 90.3%). Despite being trained on these data, AlphaFold2 failed to generate active-like activation segments for 14 kinases (**Figure 2b**). We note that multiple active-state structures are available for some kinases; therefore, averages and standard deviations are reported for those cases in **Figure 2**.

**Figure 2.**
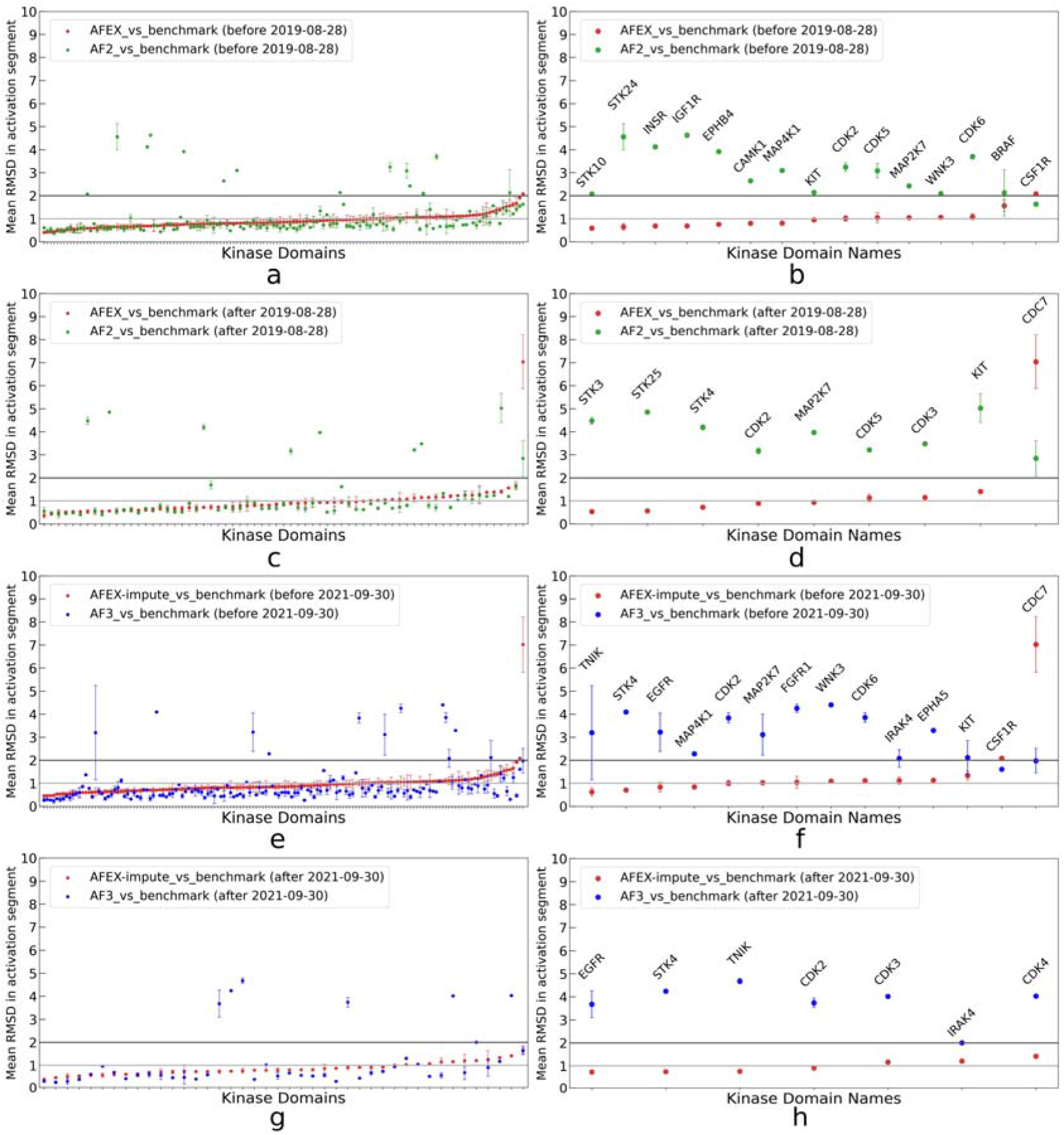
Comparison of activation segments predicted by AFEX and AlphaFold models with experimentally determined active, substrate-bound conformations of 156 different kinases from the Protein Data Bank (PDB). Panels (a)–(d) show the mean root-mean-square deviation (RMSD) and standard deviation (for kinases with multiple active structures in the PDB) of activation segments—considering all RMSD values and those exceeding 2□Å—across residues from the position preceding the DFG motif to the APE motif, evaluated using structures released before and after August 30^th^ 2019, the AlphaFold2 training cutoff date. Panels (e)–(h) present analogous comparisons for structures published before and after September 30^th^ 2021, the AlphaFold3 training cutoff. Results for AFEX and AFEX-impute are shown in red, AlphaFold2 in green, and AlphaFold3 in blue.

For 67 kinases with PDB structures of active conformation published after April 2019, AFEX maintained strong performance, achieving RMSD <1 Å in 67.1% of activation segments predictions (**Figure 2c**), with only CDC7 exceeding the 2 Å threshold (**Figure 2d**), a deviation likely due to a long disordered activation loop. AFEX models also exhibited greater consistency across datasets, maintaining approximately 99% accuracy for predictions within 2 Å in both pre- and post-April 2019 data—outperforming AlphaFold2, which achieved 92.5% accuracy and failed to generate active-like conformational states for activation segments of 9 kinases (**Figure 2d**).

To further assess AFEX’s utility in modelling ATP-complex structures, we incorporated ATP and two Mg^2^□ binding imputations using AlphaFold3, generating what we refer to as AFEX-impute models (see Methods). These models retain backbone conformations highly similar to the original AFEX predictions (<0.3□Å RMSD, **Supplementary Figure 2e**), with ATP and Mg^2^□ positioned through ligand imputation. In AlphaFold3 prediction, ATP and two Mg^2^□ ions were provided in the input, as they are essential for kinase activation. Model performance was assessed relative to AlphaFold3’s training data cutoff (September 2021), prior to which 150 active-state conformations are available. AFEX-impute models achieved activation segment predictions with RMSD <1 Å in 65.3% of cases and <2 Å in 98.7% (**and Figure 2e and 2g**). Though AlphaFold3 outperformed AFEX in the <1 Å category (80.7% vs. 65.3%), AFEX maintained superior accuracy in the <2 Å category (98.7% vs. 92.0%) (**Figure 2f**). In this range, AlphaFold3 failed to generate active-like state activation segments for 12 kinases, whereas AFEX failed only for CSF1R and CDC7. More notably, for the subset of 66 kinases with experimentally determined substrate-bound structures published after AlphaFold3’s training cutoff, AFEX achieved 71.4% <1 Å accuracy and 100% <2 Å accuracy in activation segment predictions (**Figure 2g**). AlphaFold3 matched AFEX’s <1 Å accuracy (71.4% vs. 71.4%) but underperformed at <2 Å (83.3% vs. 100.0%) (**Figure 2h**), failing to generate active-like state activation segments of 7 kinases. These results highlight AFEX’s robust performance in modeling ATP-bound kinase conformations following ATP and Mg^2^□ imputation, particularly in maintaining high accuracy in activation segment predictions within the <2 Å threshold.

Restricting the comparison to cases in which AlphaFold2 or AlphaFold3 already predicted active-like conformations (RMSD < 2 Å) showed that AFEX and AlphaFold2/3 exhibited highly similar mean RMSDs (**Supplementary Table 1**).

### Kinases with Extensive Experimental Structural Coverage

Epidermal Growth Factor Receptor (EGFR), a key member of the TYR family and a critical regulator of cellular proliferation, is extensively represented in structural databases, with 267 human EGFR structures deposited in the PDB as cataloged in KLIFS. Of these, 61 active-state conformations were determined from the 128 structures available before the AlphaFold2 training cutoff, and 78 active-state structures were resolved from the 180 structures available prior to the AlphaFold3 training cutoff. AlphaFold3 predicted inactive-like geometries even when modeled in the presence of ATP and Mg^2^□ (**Figure 3a**), whereas AFEX and AlphaFold2 successfully predicted active-state conformations of EGFR (**Figure 3b**). To elucidate the structural discrepancies among the two AlphaFold models and AFEX-impute, these three structures were superimposed, with a focused analysis on regions exhibiting substantial variation—the activation segment, C-helix, and β3 strand (**Figure 3c**). In the AlphaFold3 models, the activation loop was misoriented toward the ATP-binding pocket, sterically hindering the substrate binding site. As shown in the magnified view, one magnesium ion is slightly displaced from the ATP-binding site in the AlphaFold3-predicted structure (**Figure 3c**). In contrast, when the AFEX-predicted active-like conformation is used as a template, both magnesium ions are accurately coordinated by ATP, reflecting a functionally competent active state.

**Figure 3.**
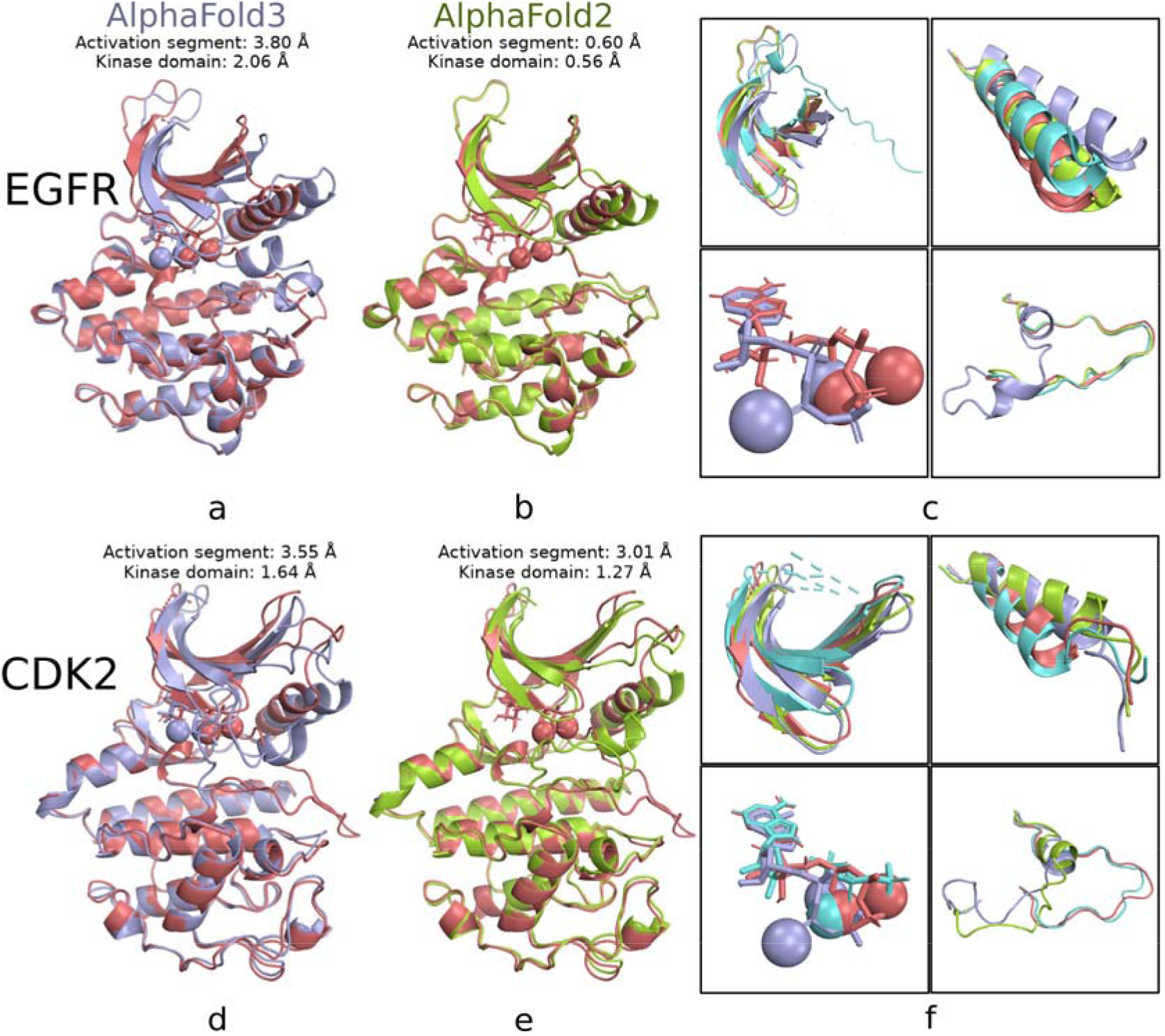
Comparison of EGFR and CDK2 structures predicted by AFEX and AlphaFold models. Panels (a) and (b) show the AFEX-impute prediction of EGFR alongside predictions from AlphaFold3 and AlphaFold2, respectively. Panels (c) displays the superimposed structures, with a detailed view of regions exhibiting significant conformational differences—the β3 strand, the C-helix, bound ligands (ATP and Mg^2+^), and the activation segment. Panel (d)-(f) present analogous comparisons for CDK2. In all panels, the AFEX-impute predictions are shown in red, the AlphaFold3 models in blue, the AlphaFold2 models in green, and representative substrate-bound PDB structures in aquamarine (EGFR: 2GS2; CDK2: 4EOM).

Cyclin-dependent kinase 2 (CDK2), a key member of the CMGC kinase family involved in cell cycle regulation, is even more extensively represented in the PDB (457 structures). Among these, 110 active-state conformations out of 284 structures were determined prior to the AlphaFold2 training cutoff date, and 115 out of 298 were resolved before the AlphaFold3 training cutoff date. Despite this substantial structural coverage, both AlphaFold2 and AlphaFold3 failed to predict the active state of CDK2, instead modeling the activation loop in substrate-blocking conformations (**Figure 3d-e**). Similar to EGFR, the AlphaFold3 prediction of CDK2 lacks a magnesium ion in the ATP-binding site, even when ATP and two magnesium ions are provided as input, indicating a deficit in functional awareness of AlphaFold3 model (**Figure 3f**). An additional example is BRAF, a representative kinase from the TKL kinase family with 109 structures deposited in the PDB, as illustrated in **Supplementary Figure 4**. Recent studies on fold-switching proteins suggest that AlphaFold models may rely more on memorized structural templates from the training data than on inference grounded in the underlying folding free energy landscape ^30^. Under this hypothesis, the presence of too many conflicting conformations in the training set may hinder AlphaFold3’s ability to resolve a consistent, functionally relevant active state.

### Consistent Prediction of Active-State Conformations in CMGC Kinases

The CMGC family comprises protein kinases involved in several critical signaling pathways, including cell cycle regulation (via cyclin-dependent kinases, CDKs), glycogen metabolism (via glycogen synthase kinase 3, GSK3), stress and growth responses (via mitogen-activated protein kinases, MAPKs), and RNA processing (via SR protein kinases). These kinases primarily catalyze the phosphorylation of serine and threonine residues and play pivotal roles in cellular proliferation, differentiation, and apoptosis, with select members also displaying tyrosine kinase activity. When we superimposed all the predicted CMGC structures together, the AFEX model - unlike the diverse and sometimes closed conformations of the activation segments predicted by AlphaFold2 and AlphaFold3 (**Figure 4a and 4b**) - revealed a distinctly open conformation of the ATP and substrate binding site (**Figure 4c**). This feature was also consistently observed across other serine/threonine kinase (STK) families upon structural superposition (**Supplementary Figure 5**).

**Figure 4.**
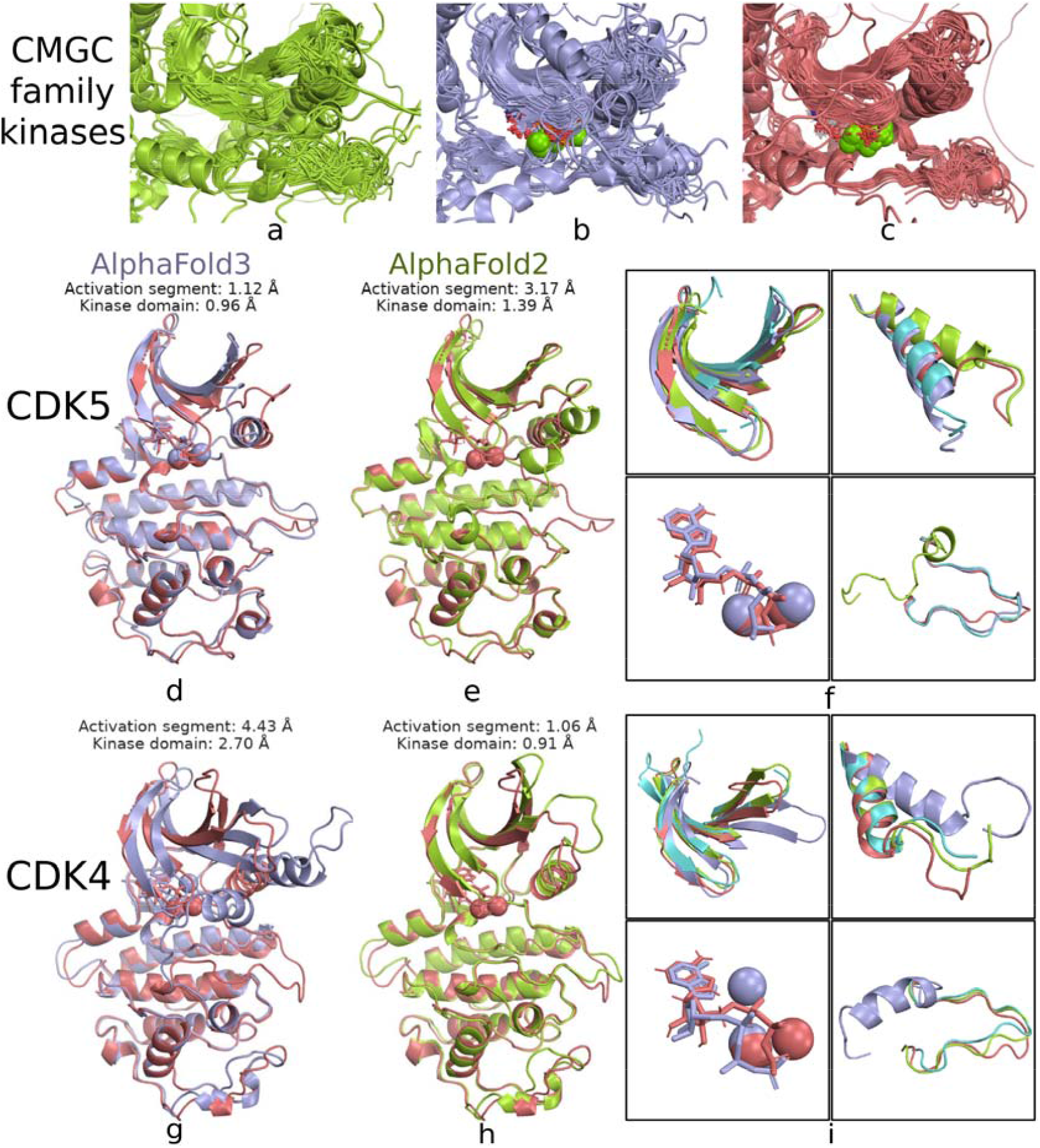
Comparison of CMGC family kinases, CDK5, and CDK4 structures predicted by AFEX and AlphaFold models. Panels (a), (b) and (c) display the superimposed structures of CMGC family kinases predicted by Alphafold2, AlphaFold3, and AFEX-impute, respectively, with focus on regions exhibiting significant conformational differences. Panels (d)-(e) show the AFEX-impute prediction of CDK5 alongside predictions from AlphaFold3 and AlphaFold2, respectively. Panels (f) presents the superimposed CDK5 structures with detailed focus on the β3 strand, the C-helix, bound ligands (ATP and Mg^2^□), and the activation segment, respectively. Panel (g)-(i) present analogous comparisons for CDK4. In all panels, the AFEX-impute predictions are shown in red, the AlphaFold3 models in blue, the AlphaFold2 models in green, and representative substrate-bound PDB structures in aquamarine (CDK5: 1UNG; CDK4: 7SJ3).

In our benchmark analysis, all CMGC kinases for which AlphaFold failed to achieve 2□Å accuracy belonged to the CDK subfamily (**Figure 2**). Both AlphaFold2 and AlphaFold3 predicted inactive conformations for CDK2, CDK3, and CDK6, despite the availability of experimentally determined active-state structures for these kinases during training. For CDK5, although the six experimentally determined structures in the PDB prior to the training cutoff all represented the active state, AlphaFold2 predicted an inactive conformation characterized by a folded activation loop, disruption of the salt bridge between the C-helix and β3 strand, and a DFG motif in the BLB-minus conformation (**Figure 4e–f**). In contrast, AlphaFold3 predicted an active-state conformation for CDK5 (**Figure 4d**), suggesting that the inclusion of ATP and Mg^2^□ in the modeling process enhanced the prediction of active kinase states—a finding also observed for BRAF (**Supplementary Figure 4d-f**) and CAMK1 (**Supplementary Figure 6a–c**), a calcium/calmodulin-dependent serine/threonine kinase from the CAMK family. Similar improvements were evident in the STK kinases from the STE family: AlphaFold2 predicted inactive conformations for STK3, STK10, STK24, and STK25, whereas AlphaFold3 successfully predicted active-state conformations when ATP and Mg^2^□ ions were included (**Supplementary Figure 7**).

However, AlphaFold3 failed to recover the active state of CDK4 in the presence of ATP and Mg^2^□ (**Figure 4g**), mispositioning the activation loop and failing to form the essential salt bridge between the C-helix and the β3 strand—a finding also observed for STK4 (**Supplementary Figure 8a–c**). In the case of CDK4, although neither model was trained on active-state structures of CDK4, AlphaFold2 successfully predicted the active conformation (**Figure 4h**), while AlphaFold3 did not (**Figure 4g and 4i**). This discrepancy highlights the stochastic nature of active-like conformation prediction when including ATP and Mg^2^□ in the input; the emergence of an open binding site is not guaranteed. A similar pattern was observed in EGFR (**Figure 3c**) and TNIK (**Supplementary Figure 8d–f**), a member of the STE kinase family, where the inclusion of ATP and Mg^2^□ resulted in a closed substrate-binding site. Interestingly, WNK3, an orphan kinase, exhibited an intermediate behavior: AlphaFold2 predicted a helical conformation in the activation loop, whereas AlphaFold3, when supplied with ATP and two magnesium ions, produced an unwound helix but lost the interaction between the N-terminal region of the activation loop and the catalytic loop, ultimately failing to predict the active state (**Supplementary Figure 9a-c**). In contrast, we highlight that AFEX-impute consistently predicted active-state conformations and generated structurally plausible coordinates for ATP and both Mg^2^□ ions across both CDKs (**Figure 4f and 4i**) and STKs (**Supplementary Figure 7 and 8c**), achieving RMSD values below 2□Å in available benchmarks (18 kinases, **Figure 2**).

### AFEX-Optimized Templates Facilitate Magnesium Ion Placement in STE Family Kinase Models

The STE kinase family was initially identified in the MAPK signaling pathway in yeast and was later shown to serve as upstream regulators in mammalian MAPK pathways. Notably, AlphaFold3 encounters difficulties in accurately modeling ATP and magnesium ion binding within the STE kinase family compared to other kinase families. Specifically, in five of the 45 catalytic domains— MAP2K3, MAP2K4, MAP2K6, MAP2K7, and TNIK—the magnesium ion is misplaced outside the ATP-binding site (**Supplementary Figure 10a**). For MAP2K4 (**Figure 5a–c**) and MAP2K6 (**Supplementary Figure 8g–i**), structurally plausible positioning of the magnesium ions was achieved only when AFEX optimized active-like structures were used as templates. This inaccuracy of ion placement in the AlphaFold models is likely due to the formation of an excessively helical activation loop, which may sterically hinder the proper coordination of the magnesium ions. A similar improvement was observed in WNK1, where the use of AFEX-predicted active-state models enabled AlphaFold3 to correctly open the ATP-binding site and position the magnesium ions accurately (**Supplementary Figure 9d-f**).

**Figure 5.**
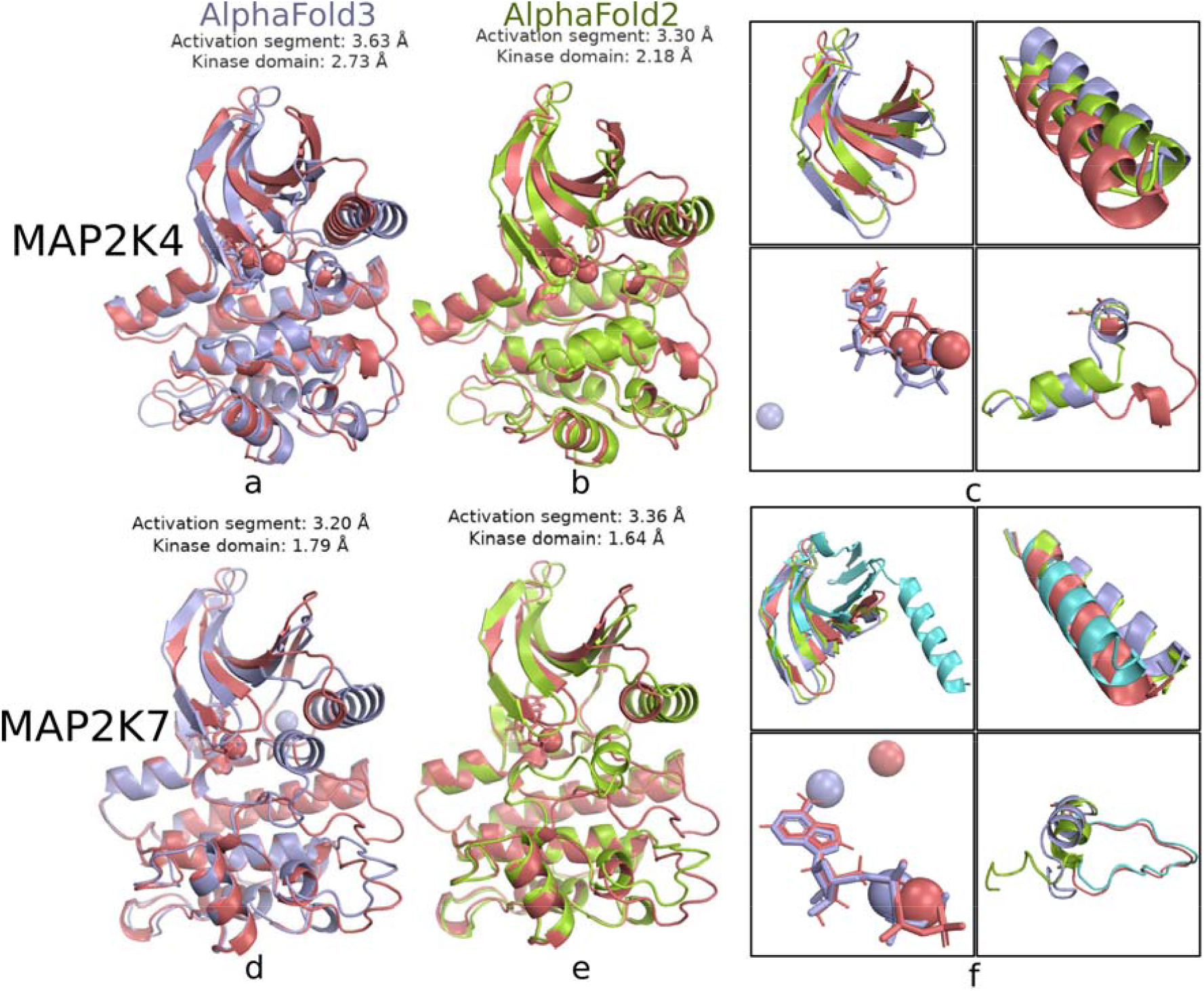
Comparison of MAP2K4 and MAP2K7 structures predicted by AFEX and AlphaFold models. Panels (a) and (b) show the AFEX-impute prediction of MAP2K4 alongside predictions from AlphaFold3 and AlphaFold2, respectively. Panels (c) displays the superimposed structures of MAP2K4, with a detailed focus on regions exhibiting significant conformational differences—the β3 strand, the C-helix, bound ligands (ATP and Mg2+), and the activation segment. Panel (d)-(f) present analogous comparisons for MAP2K7. In all panels, the AFEX-impute predictions are shown in red, the AlphaFold3 models in blue, the AlphaFold2 models in green, and representative substrate-bound PDB structures in aquamarine (MAP2K7: 6YG1).

Both AlphaFold2 and AlphaFold3 predicted inactive conformations of MAP2K7 (MKK7), characterized by a helical segment within the activation loop, the absence of a salt bridge between the C-helix and the β3 strand, and an inactive DFG motif conformation (**Figure 5d-f**). Although AFEX successfully predicted the active-state structure in agreement with the experimentally determined substrate-binding conformation, AlphaFold3 failed to accurately predict the positions of magnesium ions in MAP2K7 using the AFEX-predicted active-like structure as a template. Similar failures were observed in MAP2K3 (**Supplementary Figure 8j–l**) and TNIK (**Supplementary Figure 8d-f**), indicating persistent challenges of AlphaFold3 in predicting magnesium ion coordination within the STE kinase family.

### AFEX Consistently Predicts Open Binding Site Conformations for TYR Kinases and Rescues ATP Imputation in AATK

The TYR family comprises classical tyrosine protein kinases that catalyze the phosphorylation of tyrosine residues in proteins. This family includes both receptor tyrosine kinases, such as EGFR and INSR, and intracellular tyrosine kinases, such as SRC. The 82 catalytic kinase domains of all TYR family members were structurally aligned and compared (**Supplementary Figure 11**). Similar to observations in serine and threonine kinases, AlphaFold2 and AlphaFold3 predicted multiple closed conformations of the activation segments, whereas the AFEX structures revealed a distinct open conformation of the ATP and substrate binding site.

While some TYR kinases such as EGFR have extensive experimental structural coverage, others in this family, for example AATK, lack direct structural data in the PDB. AlphaFold2 predicts that the activation loop of AATK adopts a helical, inactive conformation (**Figures 6a**). In AlphaFold3, even with the inclusion of two magnesium ions and ATP as input, the model failed to adopt an active-like conformation; instead, ATP was mispositioned between the activation loop and the C-helix (**Figure 6a**). Only when the AFEX-predicted active-state structure was supplied as a template did AlphaFold3 successfully generate a model with plausible binding modes for both ATP and the magnesium ions (**Figure 6c**).

**Figure 6.**
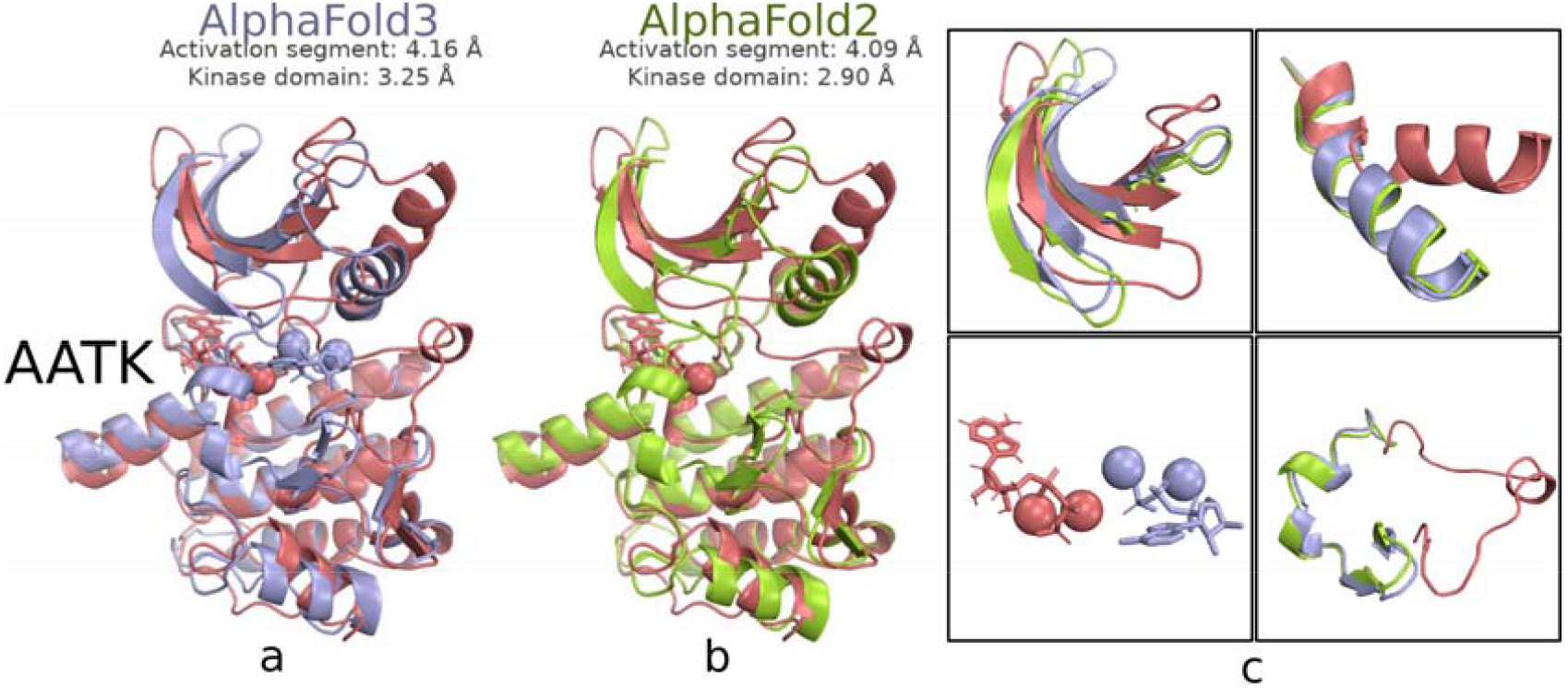
Comparison of TYR family structures and AATK structures predicted by AFEX and AlphaFold models. Panels (a) and (b) show the AFEX-impute prediction of AATK alongside predictions from AlphaFold3 and AlphaFold2, respectively. Panels (c) displays the superimposed structures of AATK, with a detailed focus on regions exhibiting significant conformational differences—the β3 strand, the C-helix, bound ligands (ATP and Mg2^+^), and the activation segment. In all panels, the AFEX-impute predictions are shown in red, the AlphaFold3 models in blue, the AlphaFold2 models in green.

## Conclusion and Discussion

In conclusion, our results demonstrate that AFEX reliably predicts active-state kinase conformations across diverse kinase families in a user-controllable manner, achieving structural accuracy sufficient to accommodate ATP and two magnesium ions within catalytic binding sites in most cases— outperforming predictions generated by AlphaFold3 alone. AFEX offers a solution to a critical limitation of current methodologies: while AlphaFold2 and AlphaFold3 often fail to generate active-state conformations, AFEX consistently produces functionally relevant geometries. A similar trend is observed with other deep-learning structure prediction methods, including ESMFold ^31^, OmegaFold ^32^, and RoseTTAFold ^33^ (**Supplementary Figure 12**). The robustness of AFEX is particularly evident in its accurate modeling of activation loops and substrate-binding regions, which are sometimes mispositioned in AlphaFold predictions, leading to steric occlusion of substrates. By preserving the functional architecture of the ATP-binding site, AFEX provides a kinome-wide framework for investigating kinase activation mechanisms and supporting structure-based design of kinase inhibitors.

This work also addresses the broader limitations of current structure prediction models in capturing conformationally dynamic protein states. Benchmark analysis revealed that AlphaFold2 and AlphaFold3 failed to predict catalytically competent conformations in a substantial number of kinases, including eight from the TYR family, eight from the STE family, five from the CMGC family, two from the TKL family, and one each from the CAMK and Other families, with RMSD values exceeding 2 Å (**Figure 2**). These mispredictions occurred even for kinases with abundant experimental active-state structural data in the PDB (**Figure 3**). Recent studies on fold-switching proteins suggest that such failures might stem from AlphaFold’s reliance on memorized structural templates rather than generalizable principles derived from the protein folding free energy landscape ^30^. Similar conclusions were drawn by Schafer et al., who showed that AlphaFold2’s improved performance over a reduced-training model stems from access to a broader repertoire of memorized templates rather than superior modeling of biophysical principles; both models displayed limited capacity to sample alternative conformations ^34^. Notably, AlphaFold often fails to accurately predict the full conformational diversity of fold-switching proteins: it typically captures only one of the experimentally determined states, with over 90% of cases missing alternative stable conformations regardless of training set inclusion. While recent work reports a ∼35% success rate for predicting fold-switching proteins likely present in the training data, AlphaFold’s ability to identify true fold switching outside of its training set remains markedly limited, as reflected by high false-negative rates and persistent bias towards one dominant conformer ^30,35^. Together, these findings highlight the potential advantage for geometry-constrained approaches like AFEX, which reduce interpolation bias and enforce biophysically meaningful constraints to improve predictive accuracy for functional states.

Our findings also show that prediction quality does not correlate with the availability of active-state training data. Although AlphaFold2 predominantly predicts active-state kinases, with approximately 70% adopting the DFG-in conformation ^14,36^, only 48% of human catalytic kinases in the EBI AlphaFold dataset exhibit catalytically competent conformations ^4^. For well-characterized kinases such as EGFR and CDK2, AlphaFold3 generated inactive-state conformations despite the abundance of structural data. The deviation of AlphaFold predictions from the geometrically defined active state in certain serine/threonine-specific kinases is unlikely to result from insufficient training data, as serine and threonine kinases possess substantially more experimentally determined active-state structures compared to tyrosine kinases ^2,37^. This observation challenges assumptions that training data abundance ensures correct functional predictions and highlights the importance of structural constraints in guiding conformational sampling. Analysis of all human CDK family kinase structures in the PDB revealed that 67% adopt inactive conformations (**Supplementary Table 2**). This imbalance in the training data likely contributes to a systematic bias toward inactive-state predictions (**Figure 3d–f** and **4d–i**). By incorporating features such as activation loop extension, DFG-in geometry, and C-helix-β3 salt bridge formation, AFEX produces diverse, family-specific active-state models that align well with experimental distributions while allowing flexibility for kinase-specific variations (**Supplementary Figure 2**). It accurately distinguishes the extended activation loops of TYR family kinases from the compact conformations of serine/threonine kinases and adapts to C-helix-β3 salt bridge variations in TKL kinases. Furthermore, AFEX accommodates atypical or orphan kinases by adapting its criteria accordingly, such as omitting salt bridge evaluation for WNK kinases lacking a β3 lysine, enabling accurate modeling across the structural diversity of the kinome.

The availability of active-state models generated by AFEX opens multiple avenues for further computational investigations. They can serve as starting points for MD simulations to study the conformational dynamics of kinase and how these dynamics are modulated by interacting ligands. The resolution of both inactive and active states is particularly useful for path-based methods, such as constant advance replicas ^38,39^ and finite-temperature string ^40^ methods. Experimental characterization of the conformational landscape of human kinases, along with the structural delineation of their distinct functional states, has already supported the validation of transition states predicted by MD and modeling studies. For example, studies on Src-family kinases (SFKs) have identified a low-populated structural intermediate that is critical for processive catalytic activity, as it regulates the rate of catalytic turnover through ADP release ^41^. AFEX-impute provides accurate structures of ATP- and Mg^2+^-bound kinase complexes across the human catalytic kinome, which could serve as suitable input structures for QM/MM simulations to probe catalytic mechanisms and chemical reactivity in a biologically relevant state. This enables mechanistic studies of activation dynamics, and facilitates kinetic modeling of signaling processes. Beyond individual kinases, these predictions may support structural inference of higher-order assemblies and complexes involving kinase domains—particularly where conformational plasticity underlies functional regulation. At the same time, although this study focuses on predicting active conformations, specific inactive conformations, particularly type-II states, are likely to be more useful for structure-based drug discovery. This will be pursued in a future study aimed at showing that AFEX-predicted structures can directly support the rational discovery of type-II kinase inhibitors.

Furthermore, the availability of functionally accurate active-state models may also facilitate the identification of kinase-substrate pairs, which is essential for elucidating cellular signaling pathways but remains a significant challenge—particularly for “dark” kinases that lack functional annotation ^42^. Traditional biochemical methods, such as affinity chromatography coupled with mass spectrometry, have been instrumental in mapping phosphorylation sites and uncovering kinase-substrate complexes ^43,44^. Yet these approaches have left over 90% of known phosphosites unassigned to a specific kinase, and nearly 30% of human kinases without any experimentally validated substrate ^42^, highlighting the scale of the knowledge gap and the need for scalable and accurate computational solutions.

While sequence-based computational methods have advanced kinase-substrate predictions ^45,46^, their performance relies heavily on curated databases such as PhosphoSitePlus ^47^ and Phospho.ELM ^48^, which are incomplete and skewed toward well-characterized kinases, thereby limiting generalizability and predictive accuracy ^49^. More importantly, these models often overlook the structural elements that determine substrate recognition. Structure-informed approaches, such as SAGEPhos ^50^, have started to incorporate 3D features to improve prediction quality. However, most available structural data lack catalytically competent kinase conformations: only a small fraction of kinase structures in public databases adopt active-like states ^4^. Integrating functional conformational states into prediction pipelines, enabled by the current work, offers a promising strategy to address these limitations. In general, by anchoring prediction models in functionally relevant structural contexts, AFEX has the potential to enhance kinase–substrate mapping and support discovery of previously uncharacterized components of signaling networks.

In summary, AFEX bridges a critical gap in current biomolecular structural prediction by enabling functionally accurate, conformation-aware representations of kinase active states. It complements existing deep learning models and provides a generalizable framework applicable to kinases across species and other protein families where functional states are defined by conserved activation signatures, including GPCRs and ion channels. While further refinement may be needed to address the dynamics of disordered loops, the integration of AFEX with other ML-based or simulation-based approaches holds promise for expanding the mechanistic toolkit of kinase biology and structure-based drug discovery.

## Methods

### Geometry-constrained Model Generation with AFEXplorer

AFEXplorer operates on the continuous MSA profile (**x**^*MSA*^) used as the actual input to the AlphaFold2 inference pipeline, rather than on the discrete MSA itself. Essentially, AFEX performs a linear transformation of the MSA feature, **wx**^*MSA*^ + **b**, with the parameter **w** and **b** iteratively updated by backpropagation through the AlphaFold2 network. The optimization is driven by a loss function that biases the output structure toward a predefined functional state while maintaining high structural plausibility. This loss function is composed of two terms (i) a CV term *L*^colvar^ that quantifies agreement with *a priori* expectation on the conformational state, and (ii) a pLDDT-based regularization term *L*^regular^ that preserves the confidence of the structural prediction. In this work, to predict the active states of catalytically competent human kinase domains, we employed five empirical geometric criteria to characterize active kinase conformations, established from systematic statistical analysis of high-confidence substrate-bound active kinase structures available in the PDB ^2,4^:

#### Active Position of the DFG Motif (DFG-in)

TThe phenylalanine (Phe) residue of the DFG motif occupies the “DFG-in” position beneath the C-helix or within a proximal pocket in the N-terminal lobe. This is quantified by two distance constraints: the Cα atom of the fourth residue following the conserved glutamate (Glu) on the C-helix (i.e., the residue that is four positions C-terminal to the Glu, denoted as Glu4-Cα) to the Cζ atom of DFG Phe (Phe-Cζ) must be less than 11 Å (d□ < 11 Å), and the Cα atom of the conserved β3 lysine (Lys-Cα) to Phe-Cζ must range between 11 and 20 Å (11 Å ≤ d□ ≤ 20 Å) ^51^.

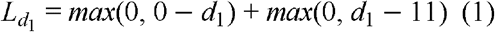

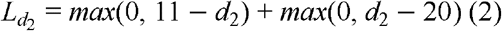

#### XDFG Dihedral Angles and BLAminus Conformation

The χ□ dihedral angle of the N, Cα, Cβ, Cγ of the Phe residue of DFG motif adopts a gauche-(g□) rotamer, corresponding to values between 240° and 360°:

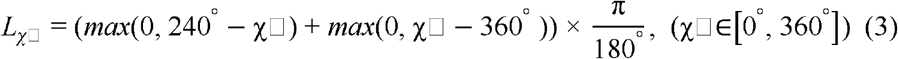

Additionally, the backbone dihedral angles (□ and ψ) of the one preceding the DFG motif (“X”), Asp, and Phe residues in the XDFG motif must align with the “BLAminus” region of the Ramachandran plot ^2^. This is assessed using cosine distance, defined for each cluster *k* in {BLAminus, ABAminus, BLBminus} as:

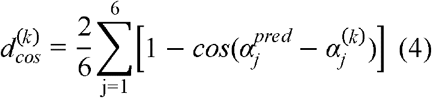

where *j* represents the six dihedral angles (□ and ψ) of the “X”, Asp, and Phe residues in the XDFG motif, respectively. 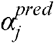 denotes the predicted value, and 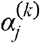 represents the corresponding predefined ideal angle for cluster *k* ^4^. The loss 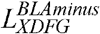 for the ‘BLAminus’ cluster is then formulated as:

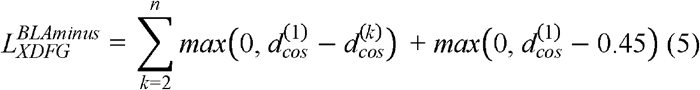

#### C-helix-β3 *Salt Bridge Distance (Salt-bridge-in)*

In the active state, a salt bridge forms between the conserved glutamate (or substitute) on the C-helix and the lysine (or substitute) in the β3 strand, stabilizing the ATP-binding site. The salt-bridge distance (d_sb_) is defined as the minimum distance between any hydrogen-bonding atoms in these residues, typically the Lys-N and Glu-O atomic distances must be less than 3.6 Å. For members of the TKL family, this threshold is relaxed to 5.1 Å. No constraint was applied when the β3 lysine is substituted to asparagine (Asn), as this substitution disrupts the salt bridge:

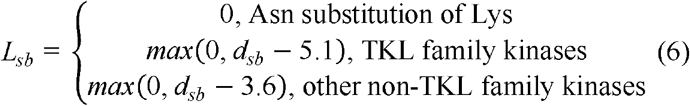

#### Activation Loop C-terminal Position (ActLoopCT-in)

The C-terminal segment of the activation loop engages the substrate in the active state. Thus, the distance between the Cα atom of the residue nine positions upstream of the APE motif (APE9-Cα) and the backbone carbonyl oxygen of the arginine in the HRD motif (Arg-O) must be less than 6.0 Å for non-tyrosine kinase (non-TYR) families and less than 8.0 Å for TYR family kinases ^4^:

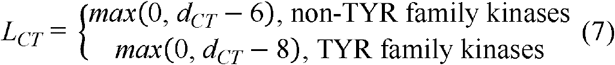

#### Activation Loop N-terminal Position (ActLoopNT-in)

The N-terminal region of the activation loop forms a backbone hydrogen bond critical for catalytic competence. This interaction occurs between the sixth residue of the DFGxxX motif (DFG6) and the residue immediately preceding the HRD motif (XHRD). The hydrogen bond is formed between the backbone amide nitrogen (N) of DFG6 and the backbone carbonyl oxygen (O) of XHRD, or *vice versa*. The minimal distance between donor and acceptor atoms (DFG6-N/O to XHRD-N/O) must be less than 3.6 Å:

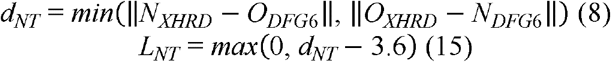

To maintain structural reliability during optimizations, we introduced a pLDDT-based regularization term that penalizes low-confidence predictions in a specified structural region:

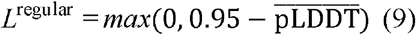

where 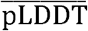 denotes the mean per-residue pLDDT value over the selected region. This regularization was computed both with respect to the catalytic domain, 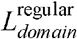, and to the activation loop segment, 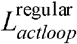. The penalty is zero when the mean pLDDT of the corresponding region is at least 0.95 and increases linearly as confidence decreases below this threshold.

The total loss function for AFEX optimization is the weighted sum of all geometric and confidence-based terms. These geometric criteria are added together to form the following loss function for AFEX optimization:

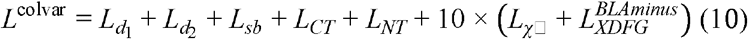

The total AFEX optimization loss is then:

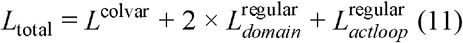

Models were deemed optimized when all five geometric criteria were satisfied and the average pLDDT score exceeded 0.95 (i.e., *L*_total_ = 0). Although 50 refinement iterations were executed, final models were selected from the 25^th^ iteration onward to ensure convergence, with RMSD between the 25^th^ and 50^th^ iteration structures consistently below 0.1 Å. We also note that the following exceptions to geometric criteria were considered for certain kinases due to their unique structural features:

#### APE Motif Variants

Kinases such as BUB1, HASPIN, TP53RK, and PKDCC lacked a canonical APE motif, resulting in altered C-terminal activation loop architecture. These were excluded from ActLoopCT-in requirements.

#### Long Disordered Activation Loop

MASTL kinase was excluded due to an exceptionally long and highly disordered activation loop (575 amino acids), which precluded reliable structural modelling within the AFEX framework.

#### Computational cost

AFEX incurs additional computational cost due to iterative optimization during inference, requiring approximately 10 minutes to 4 hours per kinase to generate active-state conformations on a single NVIDIA A40 GPU.

### Kinase Structure Prediction Using AlphaFold

The structures of human protein kinase catalytic domains were predicted using AlphaFold2 ^16^ and AlphaFold3 ^17^, developed by Google DeepMind. For AlphaFold2 predictions, the monomer model was used. The FASTA formatted amino acid sequences for kinase catalytic domains were obtained from the Kincore database ^4^ and used for input. When predictions were conducted in template-disabled mode in AlphaFold2, the “--max_template_date” was set to 1700-01-01 with a reduced database preset (--db_preset reduced_dbs). Precomputed MSA function was enabled (--use_precomputed_msas), and all other parameters were kept at their default values. For AlphaFold3 predictions, input amino acid sequences were provided in JSON format.

### Imputation of ATP and Mg^2+^ Ions

Ligands and ions were subsequently imputed into the active-state kinase domains by aligning the AlphaFold3-predicted complexes with the AFEX models for *in situ* imputation. The kinase structures containing ligands and ions underwent default structural energy minimisation using Chimera 1.18 ^52^ to eliminate potential atomic clashes. Key residues critical for active-state conformations were frozen during minimisation to preserve structural integrity. These residues included the XHRD and XDFG motifs, Glu and Glu4 on the C-helix, β3 lysine, APE9, and DFG6. Before energy minimisation, the AFEX-impute backbone is exactly the same as the original AFEX backbone. Both pre- and post-imputed AFEX kinase structures were confirmed to maintain active-state conformations as assessed by Kincore ^14^, with RMSD values of less than 0.3□Å relative to the original AFEX active-state model (**Supplementary Figure 2e**).

### Structure Prediction Using ESMFold, OmegaFold, and RoseTTAFold

The structures of human protein kinase catalytic domains were predicted using ESMFold (esmfold_v1) ^31^ and OmegaFold ^32^, installed and executed according to the instructions provided in their respective GitHub repositories. The all-atom version of RoseTTAFold ^33^ was used to predict kinase-ATP-Mg^2+^ complexes. Sequence databases including UniRef30 (2020_06), BFD (first_non_consensus_sequences), PDB100 (2021Mar03), and BLAST (v2.2.26) were employed for MSA and template generation. Input amino acid sequences were provided as FASTA files, and ATP and Mg^2+^ ions were specified as SMILES strings within YAML input files.

### Benchmark Test

The predicted active-state models were validated by comparing them against 156 PDB structures that met substrate-binding activity criteria. Validation relied on root-mean-square deviation (RMSD) calculations to measure the average atomic displacement between predicted and experimentally resolved structures. The alignment and RMSD computation were conducted using PyMOL’s automated multi-step superposition algorithm. This algorithm employs sequence alignment based on BLOSUM62 weightings ^53^ and only one cycle of iterative refinement to reject outliers, focusing on backbone atoms (N, Cα, C, O). After aligning the model and reference structures, RMSD values for the activation segment backbone atoms were calculated using the “rms_cur” function in PyMOL, which computes RMSD without performing additional fitting.

## Data Availability and Reproducibility

The initial input comprises feature.pkl files generated by AlphaFold2 for each target sequence. FASTA files for the 436 human catalytic protein kinase domains used to generate these features are available from the Kincore database at https://dunbrack.fccc.edu/kincore/static/downloads/af2activemodels/kinasedomainfasta.tar.gz. The final output is a PDB file representing the predicted kinase structure that satisfies the specified active-state criteria. The AFEX active-state structural models for all 436 catalytic kinases, including those with imputed ATP- and magnesium-bound states, are publicly accessible at https://github.com/JingHuangLab/AFEX-kinome/tree/main/database. The benchmarking dataset comprising 3,010 experimentally determined active structures of 156 human catalytic protein kinases from the PDB used in this study is available at https://github.com/JingHuangLab/AFEX-kinome/releases/tag/benchmark-2025-10-23.

## Author Contributions

JH and YW conceived and designed the study. YW performed coding, structure prediction, and data analysis. JH, YW, and QH drafted the manuscript, and all authors contributed to manuscript revision and approved the final version.

## Acknowledgments

We thank Dr. Zilin Song, Dr. Tengyu Xie, Dr. Jing Xue, and Ms. Runtong Qian for insightful discussions. This work is supported by the Shenzhen Medical Research Fund (B2404003), the National Natural Science Foundation of China (T2596084), the “Pioneer” and “Leading Goose” R&D Program of Zhejiang (2023C03109), the Westlake University-Muyuan Joint Research Institute (WU2025MY001), the State Key Laboratory of Gene Expression, and the Westlake Education Foundation. We thank the Westlake University Supercomputer Center for computational resources and related assistance.

## Supplemental information

Document S1. Supplementary Figures 1–12, Supplementary Tables 1 and 2

