## Supporting Information for "Geometry-Constrained Prediction of Catalytically Competent Kinase Domains Across the Human Kinome"

### Supporting Information for
### Geometry-Constrained Prediction of Catalytically Competent Kinase Domains Across the Human Kinome

**Yuxuan Wang [1,2,3], Qi Hu [1,2,3], Jing Huang [1,2,3,*]**
1 State Key Laboratory of Gene Expression, School of Life Sciences, Westlake University, Hangzhou 310030, Zhejiang, China
2 Westlake AI Therapeutics Lab, Westlake Laboratory of Life Sciences and Biomedicine, Hangzhou 310024, Zhejiang, China
3 Institute of Biology, Westlake Institute for Advanced Study, Hangzhou 310024, Zhejiang, China

#### Supplementary Figures

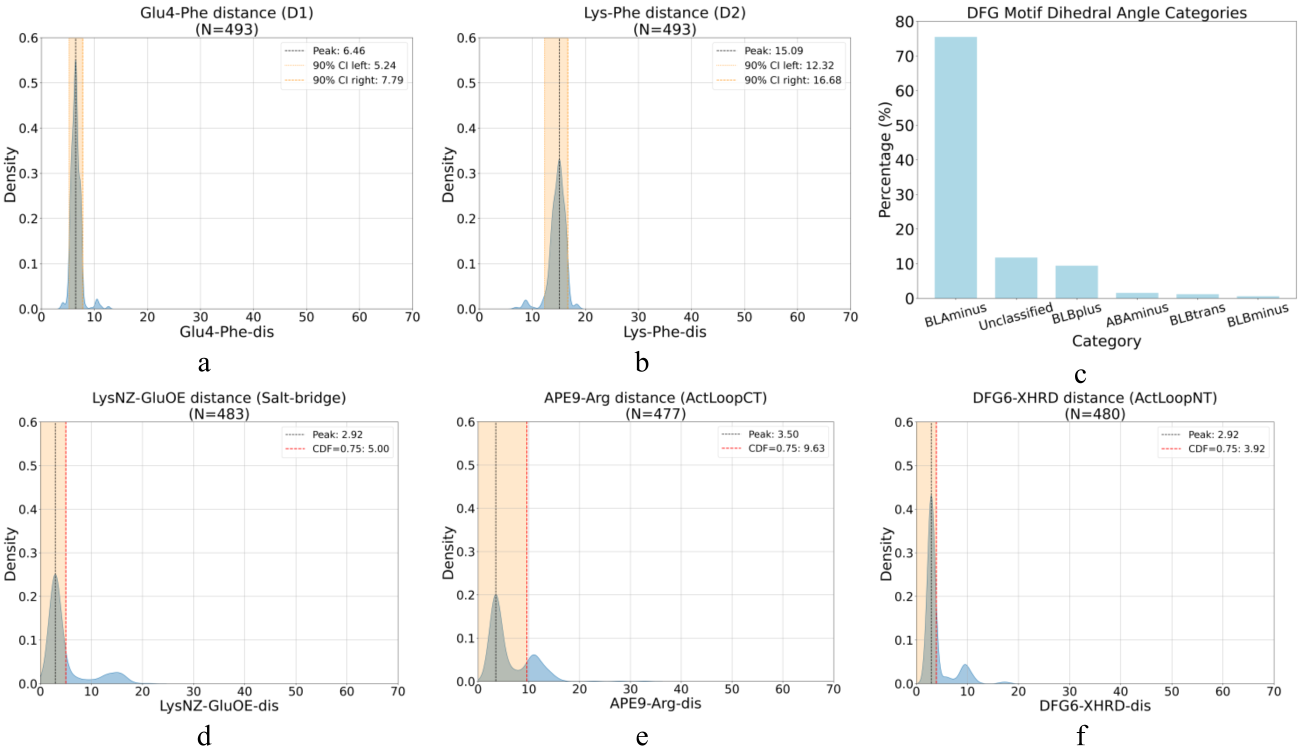


**Supplementary Figure 1. Statistical analysis of key geometric parameters in ATP/ADP-bound kinase structures from the PDB.** (a, b) The distance between the Glu4-Cα and the Phe-Cζ (D1), and between the Lys-Cα and the Phe-Cζ (D2), consistent with a DFG-in conformation. (c) Dihedral angle analysis reveals that 75.5% of ATP/ADP-bound kinase structures in the PDB adopt the BLAminus conformation in the DFG motif. (d) The distance distribution between the Oε atom of the C-helix glutamate and the Nζ atom of the β3 lysine shows a peak at 2.92 Å. (e) The distance between the Cα atom of the residue nine positions upstream of the APE motif (APE9-Cα) and the HRD arginine peaks at 3.50 Å, with a cumulative density of 0.75 extending to 9.63 Å, suggesting interactions between the activation loop and catalytic loop. (f) The distance between the backbone atoms of the sixth residue in the DFGxxX motif (DFG6) and the residue preceding the HRD motif (XHRD) peaks at 2.92 Å, with a cumulative density of 0.75 at 3.92 Å, supporting the presence of a likely backbone–backbone hydrogen bond. Key geometric thresholds were derived from Roland et al., (see Methods).

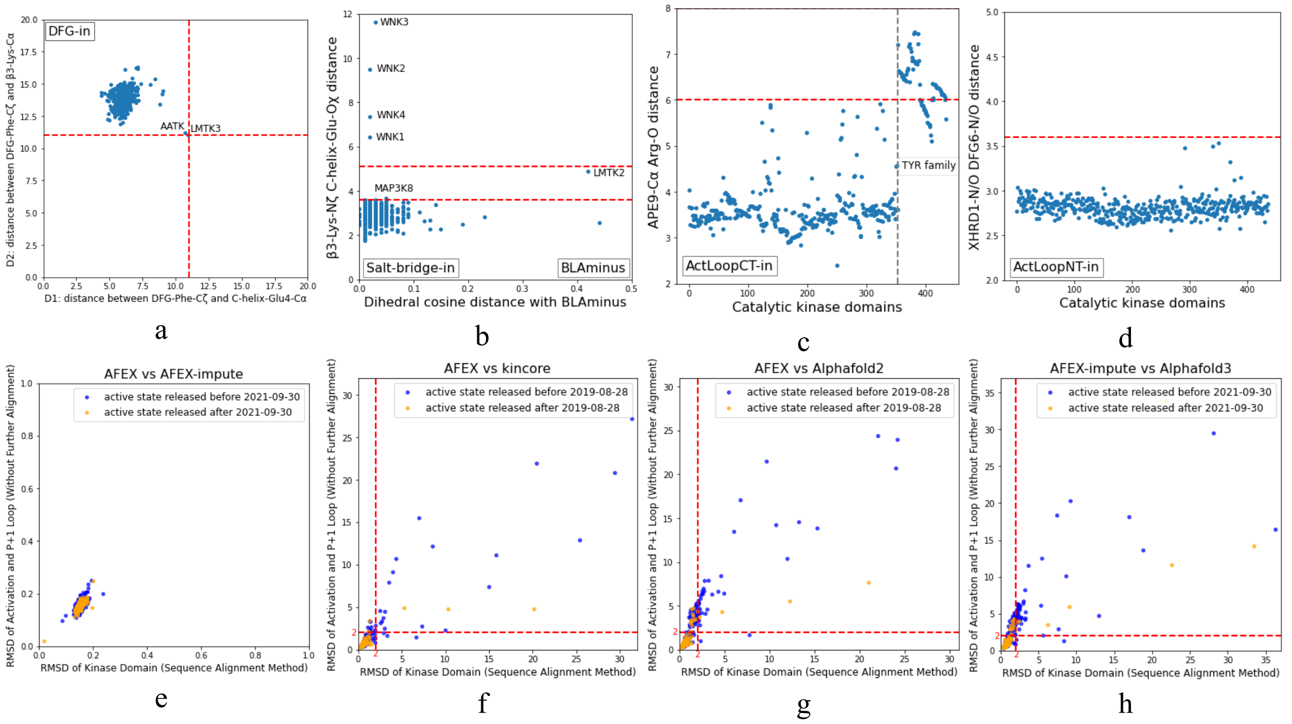


**Supplementary Figure 2. Structural geometry analysis and comparative accuracy assessment of AFEXplorer-predicted active-state conformations across 436 human kinase catalytic domains.** (a) Spatial distribution of phenylalanine residues within the DFG motif across all kinase domains. (b) Salt bridge distances within the N-terminal domain, paired with cosine distances to the BLAminus conformation in the Ramachandran plot. (c) Activation segment geometry focusing on the distance between the Cα atom of the residue nine positions upstream of the APE motif (APE9-Cα) and the backbone carbonyl oxygen of the arginine in the HRD motif (Arg-O). (d) N-terminal positioning of the P+1 loop relative to the catalytic cleft, focusing on the interaction between the sixth residue in the DFGxxX motif (DFG6) and the residue immediately preceding the HRD motif. (e) Structural divergence in RMSD between AFEX structures and AFEX-impute structures which complexed with AlphaFold3-derived ATP and $Mg^{2+}$ ions following energy minimization (see Methods). (f) Structural divergence between AFEX structures against the Kincore dataset. (g) Structural divergence between AFEX-predicted active-state conformations and AlphaFold2-generated models. (h) Structural divergence between AFEX-impute models and standard AlphaFold3 predictions incorporating ATP and Mg2+ ions. Orange and blue markers distinguish whether experimentally resolved kinase structures can be retrieved before and after the specified cut-off date, respectively.

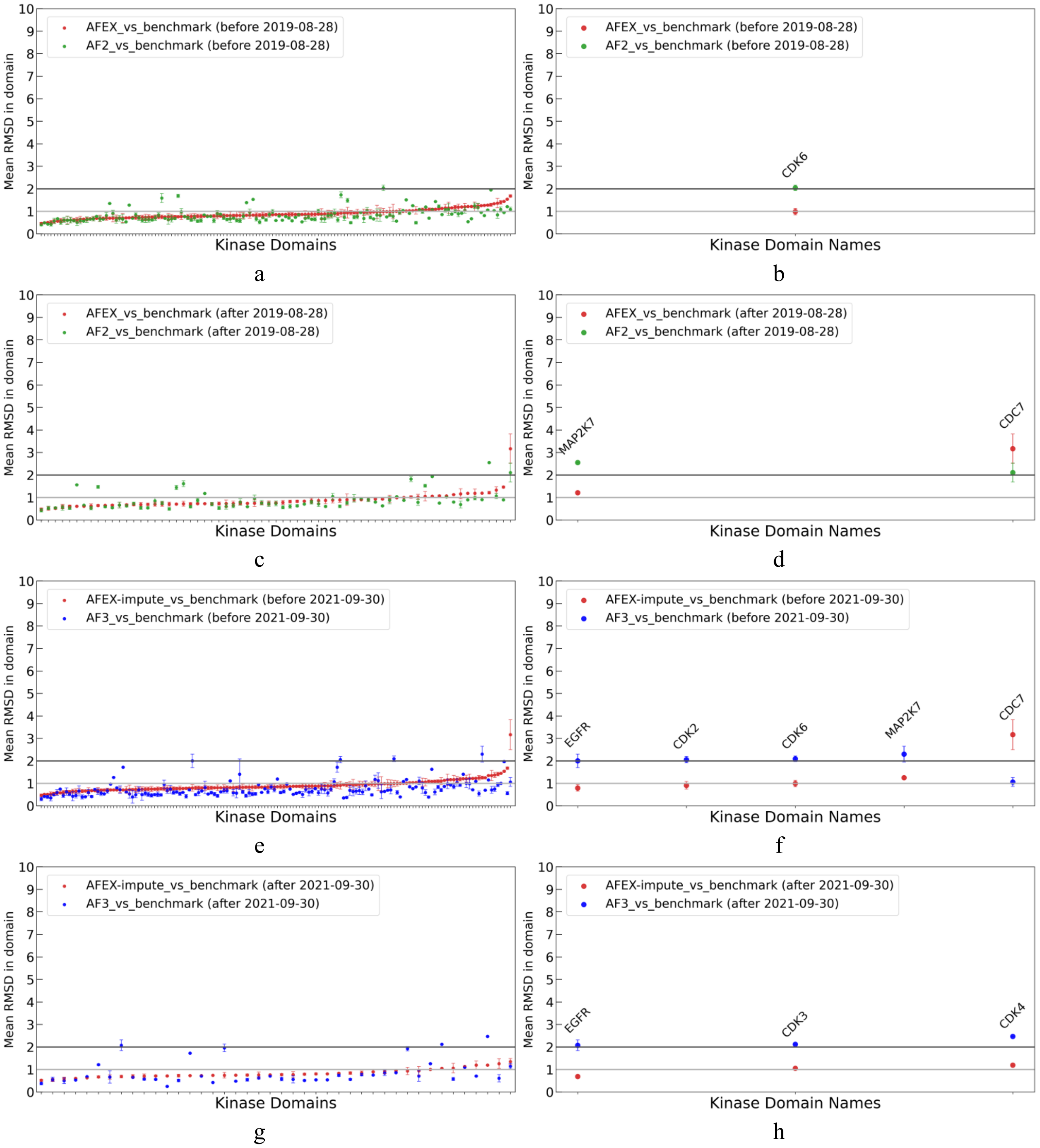


**Supplementary Figure 3. Comparison of catalytic kinase domains predicted by AFEX and AlphaFold models with experimentally determined active, substrate-bound conformations of 156 kinases from PDB.** Panels (a)–(d) display the mean RMSD and standard deviation (for kinases with multiple active structures in the PDB), including all RMSD values and those exceeding 2 Å, calculated across all residues within kinase domains. Analyses are stratified by structures released before and after August 30$^{th}$, 2019, the AlphaFold2 training cutoff date. Panels (e)–(h) present analogous comparisons for structures published before and after September 30$^{th}$, 2021, the AlphaFold3 training cutoff. Results for AFEX and AFEX-impute are shown in red, AlphaFold2 in green, and AlphaFold3 in blue.

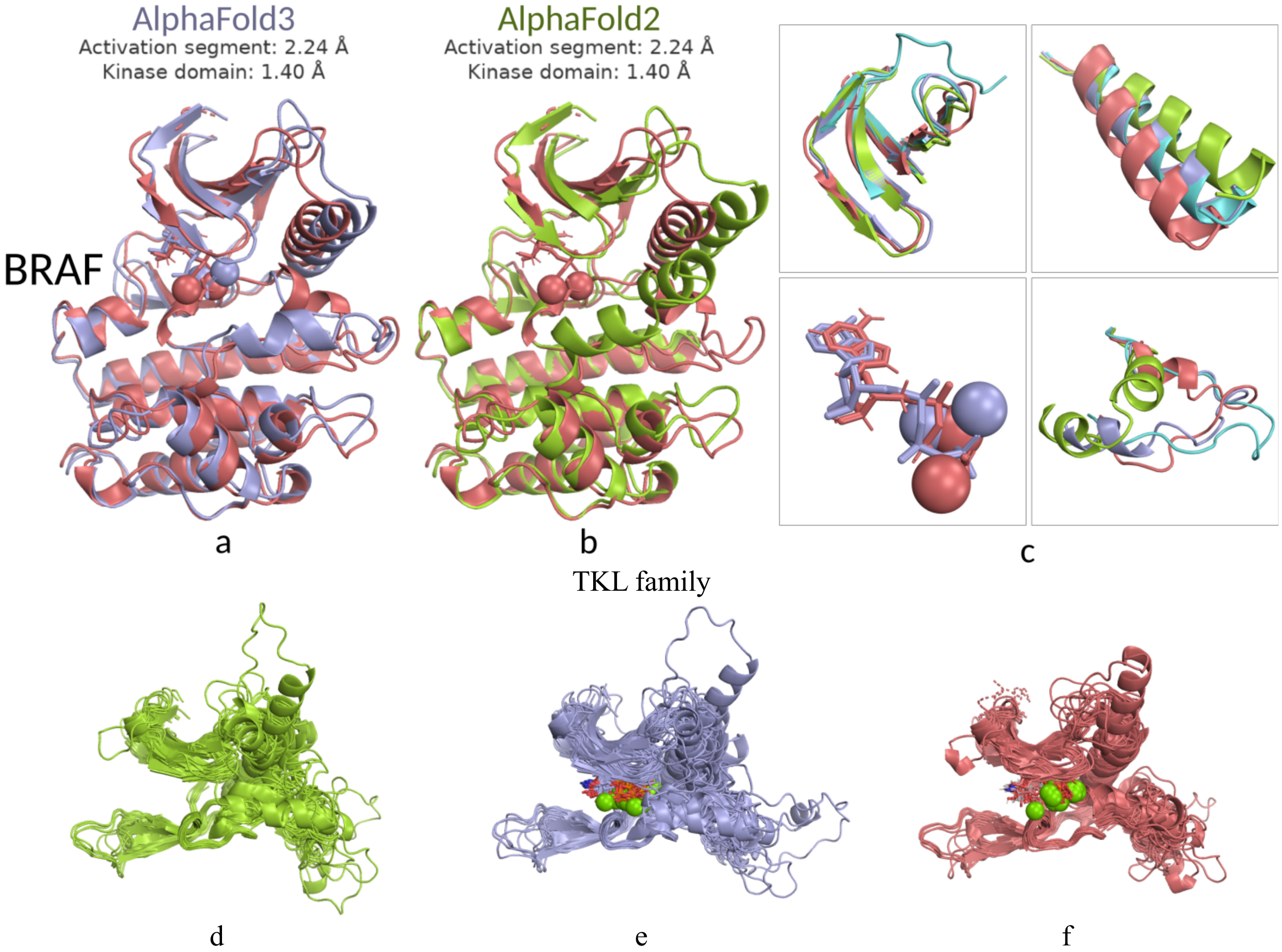


**Supplementary Figure 4. Comparison of TKL family structures and BRAF structures predicted by AFEX and AlphaFold models.** Panels (a) and (b) show the AFEX-impute prediction of BRAF alongside predictions from AlphaFold3 and AlphaFold2, respectively. Panels (c) displays the superimposed structures of BRAF, with a detailed focus on regions exhibiting significant conformational differences—the β3 strand, the C-helix, bound ligands (ATP and $Mg^{2+}$), and the activation segment. Panels (d), (e) and (f) display the superimposed structures of TKL family kinases predicted by Alphafold2, AlphaFold3, and AFEX-impute, respectively, with focus on regions exhibiting significant conformational differences. In all panels, the AFEX-impute predictions are shown in red, the AlphaFold3 models in blue, the AlphaFold2 models in green, and representative substrate-bound PDB structures in aquamarine (BRAF: 4MNE).

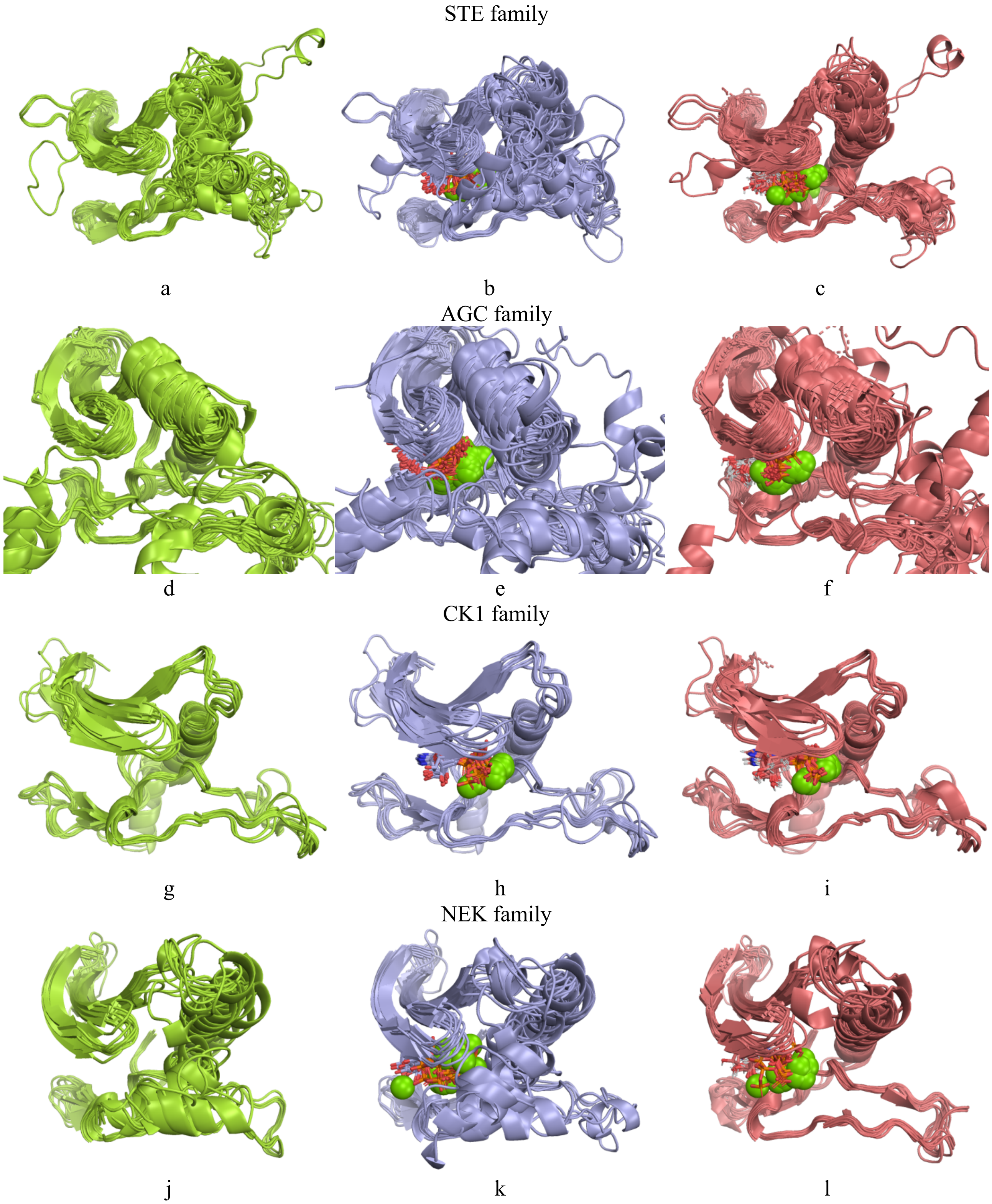


**Supplementary Figure 5. Superimposed structures of STE family kinases, AGC family kinases, CK1 family kinases, and NEK family kinases predicted by AFEX and AlphaFold models.** Panels (a), (b) and (c) display the superimposed structures of STE family kinases predicted by Alphafold2, AlphaFold3, and AFEX-impute, respectively, with a detailed focus on regions exhibiting significant conformational differences—the activation segment, C-helix, and β3 strand. Panels (d)–(f) present corresponding comparisons for AGC family kinases, (g)–(i) for CK1 family kinases, and (j)–(l) for NEK family kinases. In all panels, AFEX-impute predictions are shown in red, AlphaFold3 models in blue, AlphaFold2 models in green.

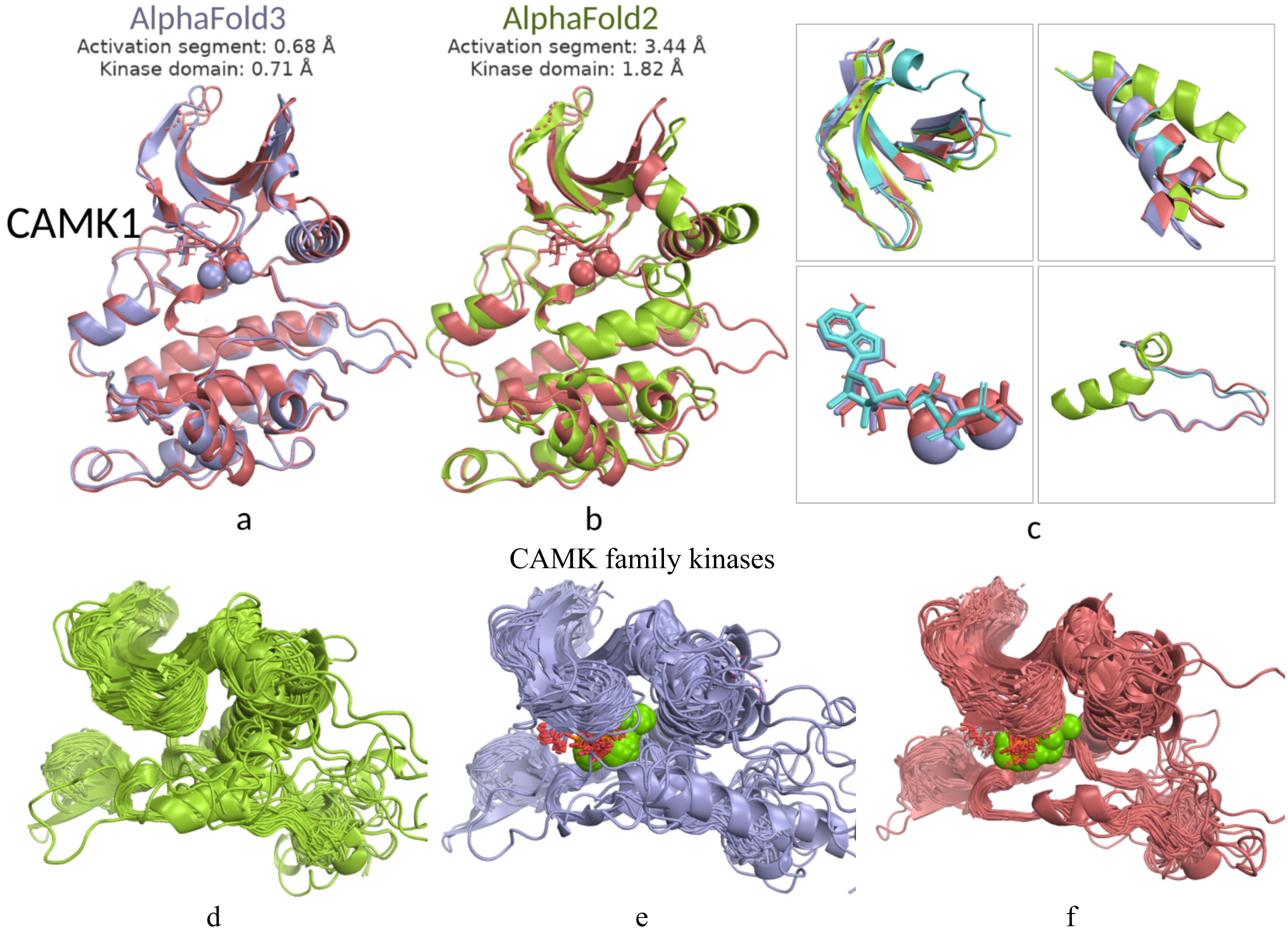


**Supplementary Figure 6. Comparison of CAMK family structures and CAMK1 structures predicted by AFEX and AlphaFold models.** Panels (a) and (b) show the AFEX-impute prediction of CAMK1 alongside predictions from AlphaFold3 and AlphaFold2, respectively. Panels (c) displays the superimposed structures of CAMK1, with a detailed focus on regions exhibiting significant conformational differences—the β3 strand, the C-helix, bound ligands (ATP and $Mg^{2+}$), and the activation segment. Panels (d), (e) and (f) display the superimposed structures of CAMK family kinases predicted by Alphafold2, AlphaFold3, and AFEX-impute, respectively, with focus on regions exhibiting significant conformational differences. In all panels, the AFEX-impute predictions are shown in red, the AlphaFold3 models in blue, the AlphaFold2 models in green, and representative substrate-bound PDB structures in aquamarine (CAMK1: 4FG7).

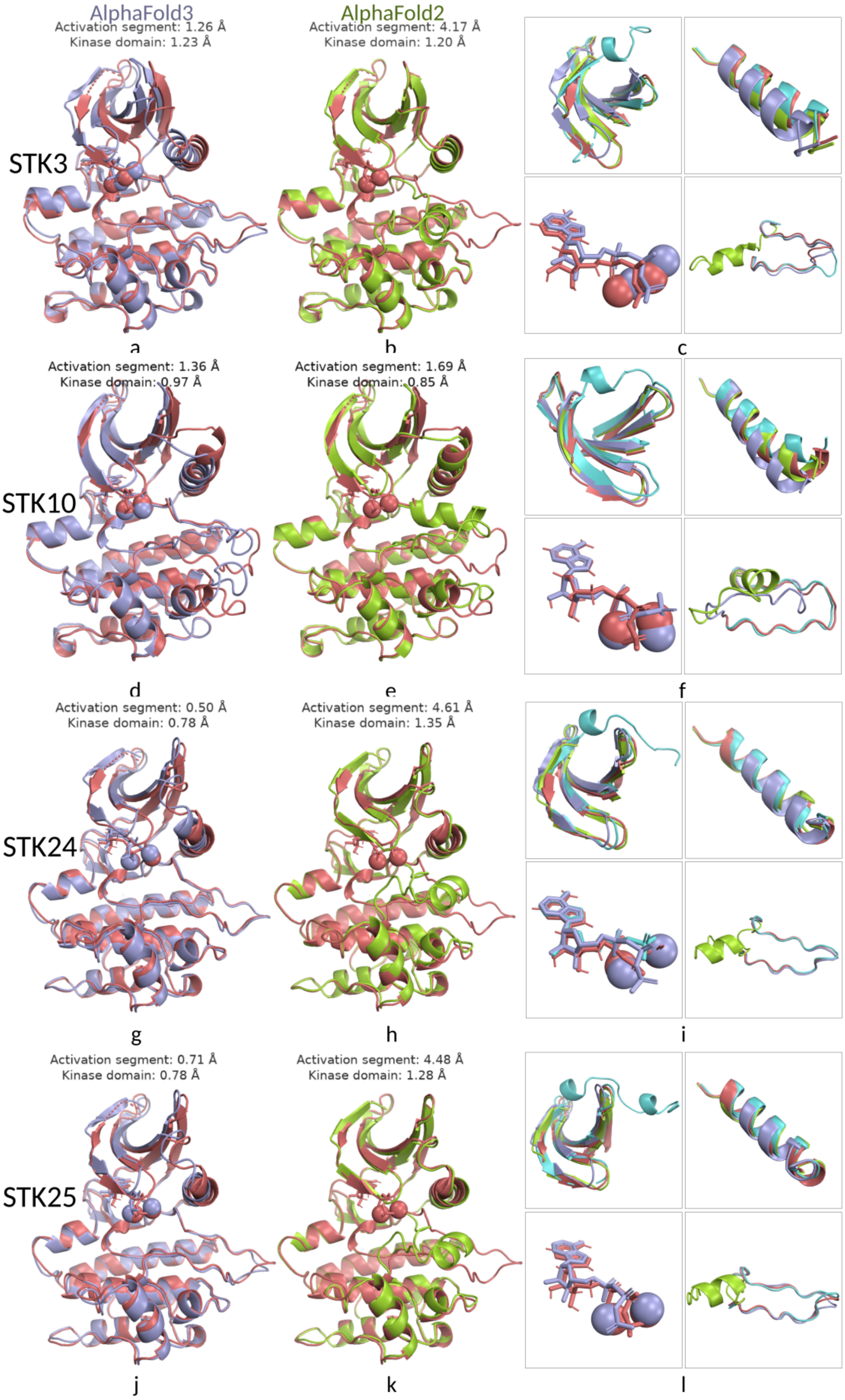


**Supplementary Figure 7. Structural comparison of STK3, STK10, STK42, and STK25 predicted by AFEX and AlphaFold models.** Panels (a) and (b) display the AFEX-impute prediction of STK3 alongside predictions from AlphaFold3 and AlphaFold2, respectively. Panel (c) shows the superimposed structures of STK3, highlighting regions with notable conformational differences—the activation segment, C-helix, and β3 strand. Panels (d)–(f) present corresponding comparisons for MAP2K7, (g)–(i) for STK24, and (j)–(l) for STK25. In all panels, AFEX-impute predictions are shown in red, AlphaFold3 models in blue, AlphaFold2 models in green, and representative substrate-bound PDB structures in aquamarine (STK3: 8A66; STK10: 6HXF; STK24: 3A7J; STK25: 7Z4V).

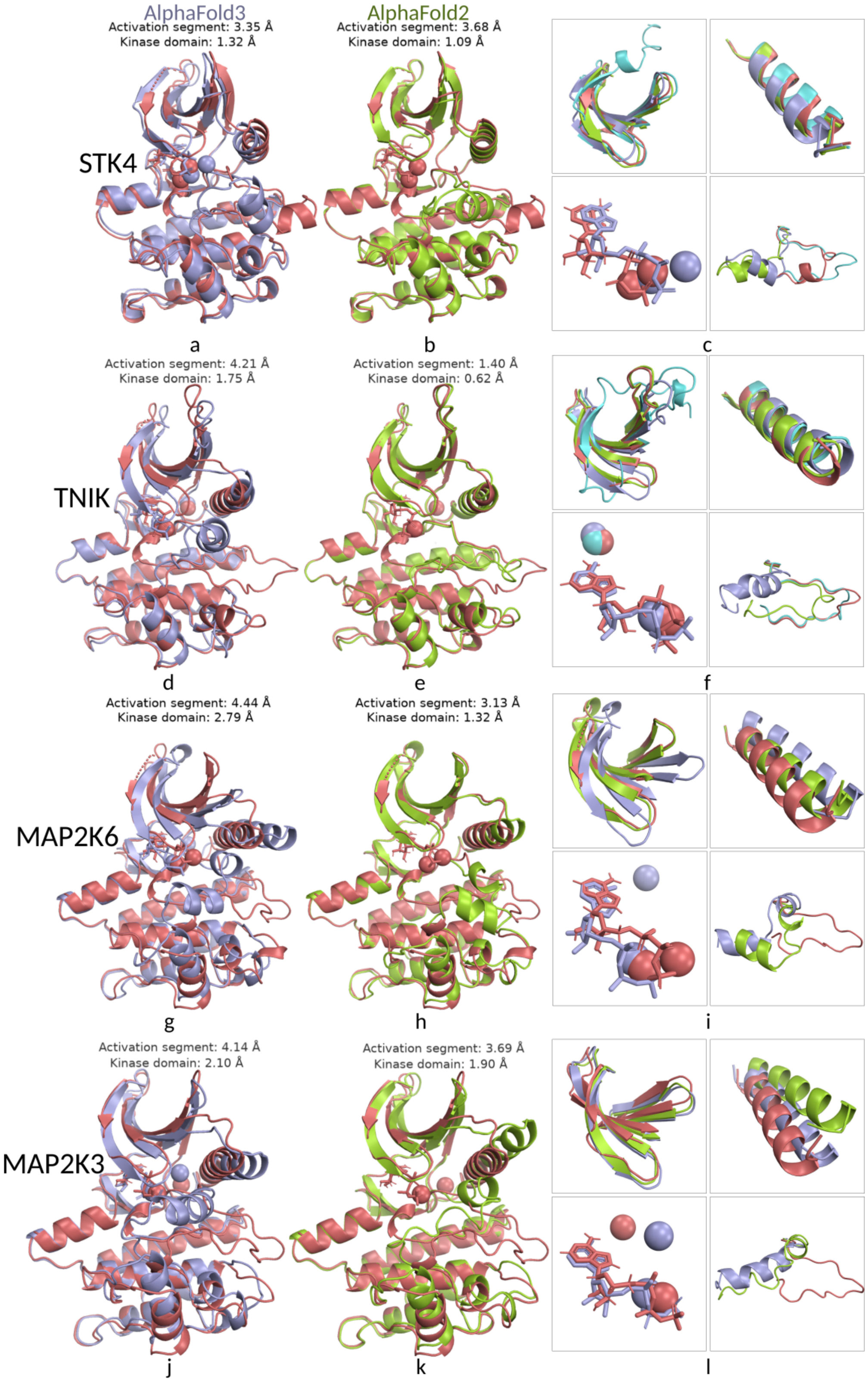


**Supplementary Figure 8. Structural comparison of STK4, TNIK, MAP2K6, and MAP2K3 predicted by AFEX and AlphaFold models.** Panels (a) and (b) show the AFEX-impute prediction of STK4 alongside predictions from AlphaFold3 and AlphaFold2, respectively. Panels (c) displays the superimposed structures of STK4, with a detailed focus on regions exhibiting significant conformational differences—the β3 strand, the C-helix, bound ligands (ATP and $Mg^{2+}$), and the activation segment. Panels (d)–(f) present corresponding comparisons for TNIK, (g)–(i) for MAP2K6, and (j)–(l) for MAP2K3. In all panels, AFEX-impute predictions are shown in red, AlphaFold3 models in blue, AlphaFold2 models in green, and representative substrate-bound PDB structures in aquamarine (STK4: 6YAT; TNIK: 6RA7).

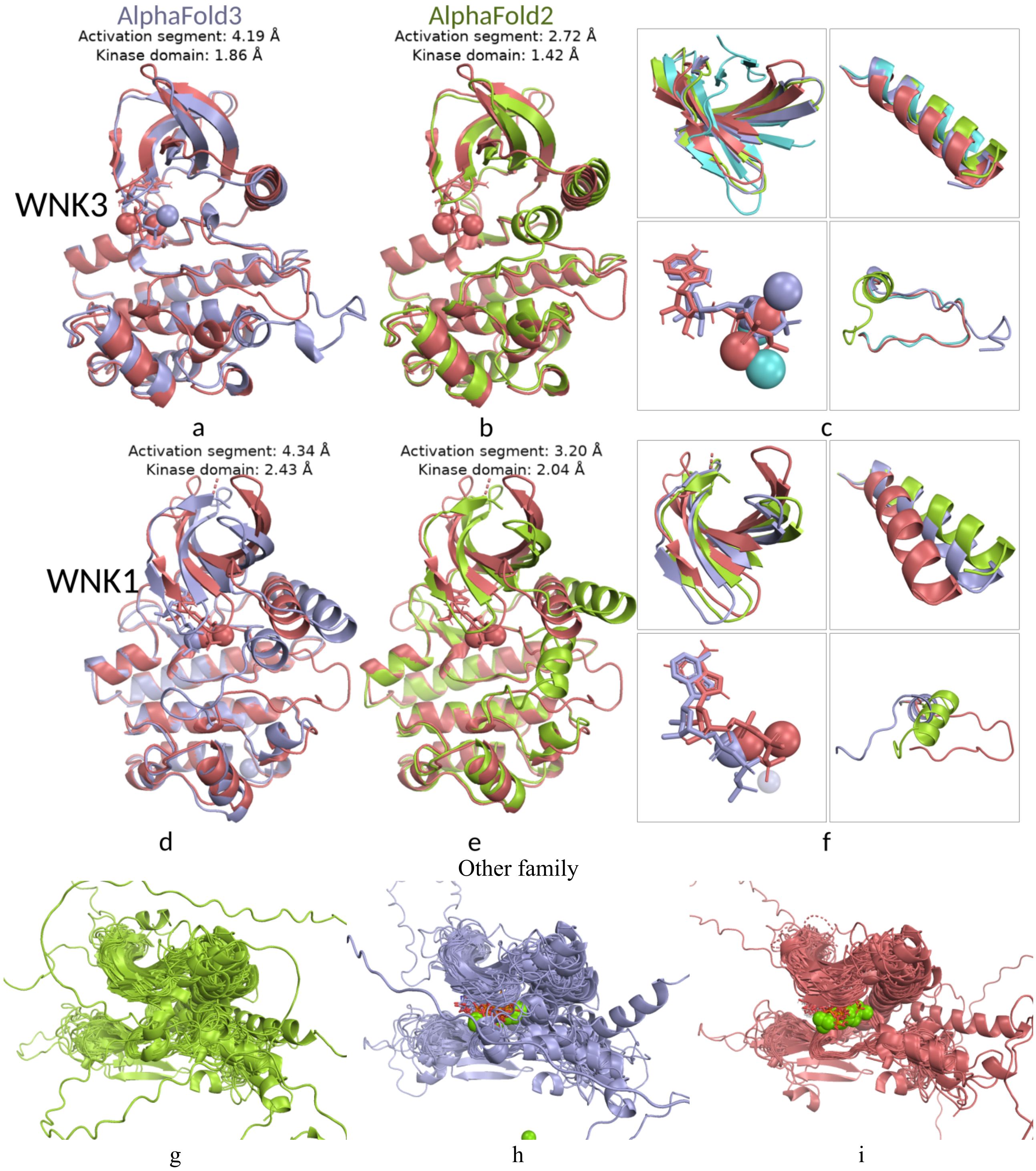


**Supplementary Figure 9. Comparison of Other family structures, WNK3 structures, and WNK1 structures predicted by AFEX and AlphaFold models.** Panels (a) and (b) show the AFEX-impute prediction of WNK3 alongside predictions from AlphaFold3 and AlphaFold2, respectively. Panels (c) displays the superimposed structures of WNK3, with a detailed focus on regions exhibiting significant conformational differences—the β3 strand, the C-helix, bound ligands (ATP and $Mg^{2+}$), and the activation segment. Panels (d)–(f) present corresponding comparisons for WNK1. Panels (g), (h) and (i) display the superimposed structures of Other family kinases predicted by Alphafold2, AlphaFold3, and AFEX-impute, respectively, with focus on regions exhibiting significant conformational differences. In all panels, the AFEX-impute predictions are shown in red, the AlphaFold3 models in blue, the AlphaFold2 models in green, and representative substrate-bound PDB structures in aquamarine (WNK3: 5O26).

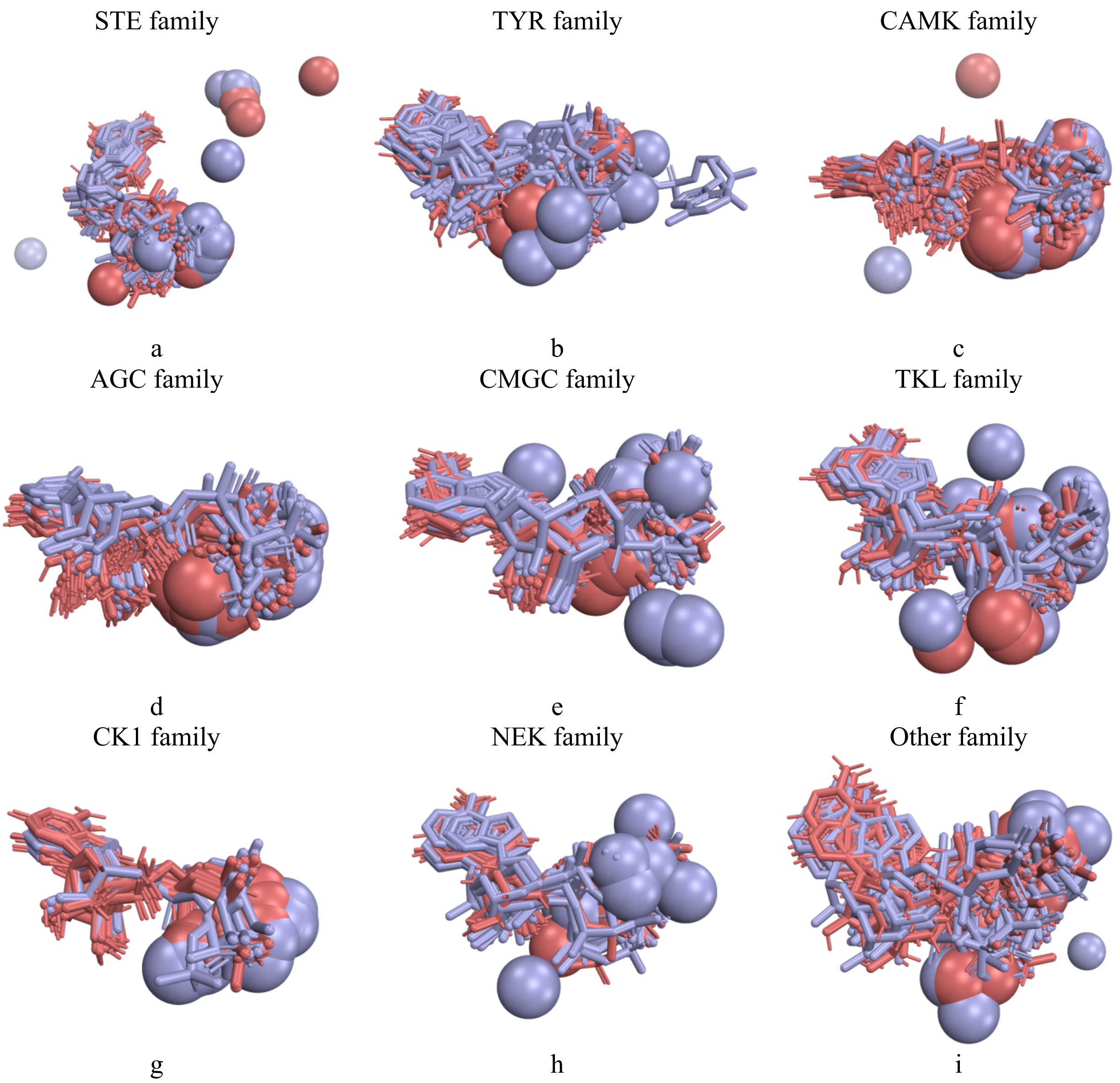


**Supplementary Figure 10. Superimposition of ATP and two magnesium ions across each kinase family.** AFEX-impute results were shown in red and AlphaFold3 predictions in blue. For each plot, ATP molecules and magnesium ions were extracted and superimposed by aligning the original structures to a shared reference structure from the corresponding kinase family.

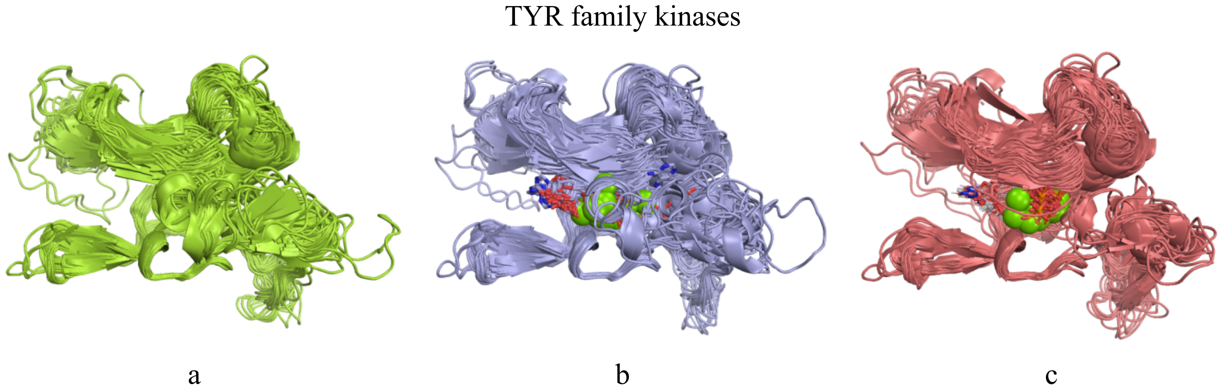


**Supplementary Figure 11. Comparison of TYR family structures predicted by AFEX and AlphaFold models.** Panels (a), (b) and (c) display the superimposed structures of TYR family kinases predicted by Alphafold2, AlphaFold3, and AFEX-impute, respectively, with focus on regions exhibiting significant conformational differences—the activation segment, C-helix, and β3 strand. The AFEX-impute predictions are shown in red, the AlphaFold3 models in blue, the AlphaFold2 models in green.

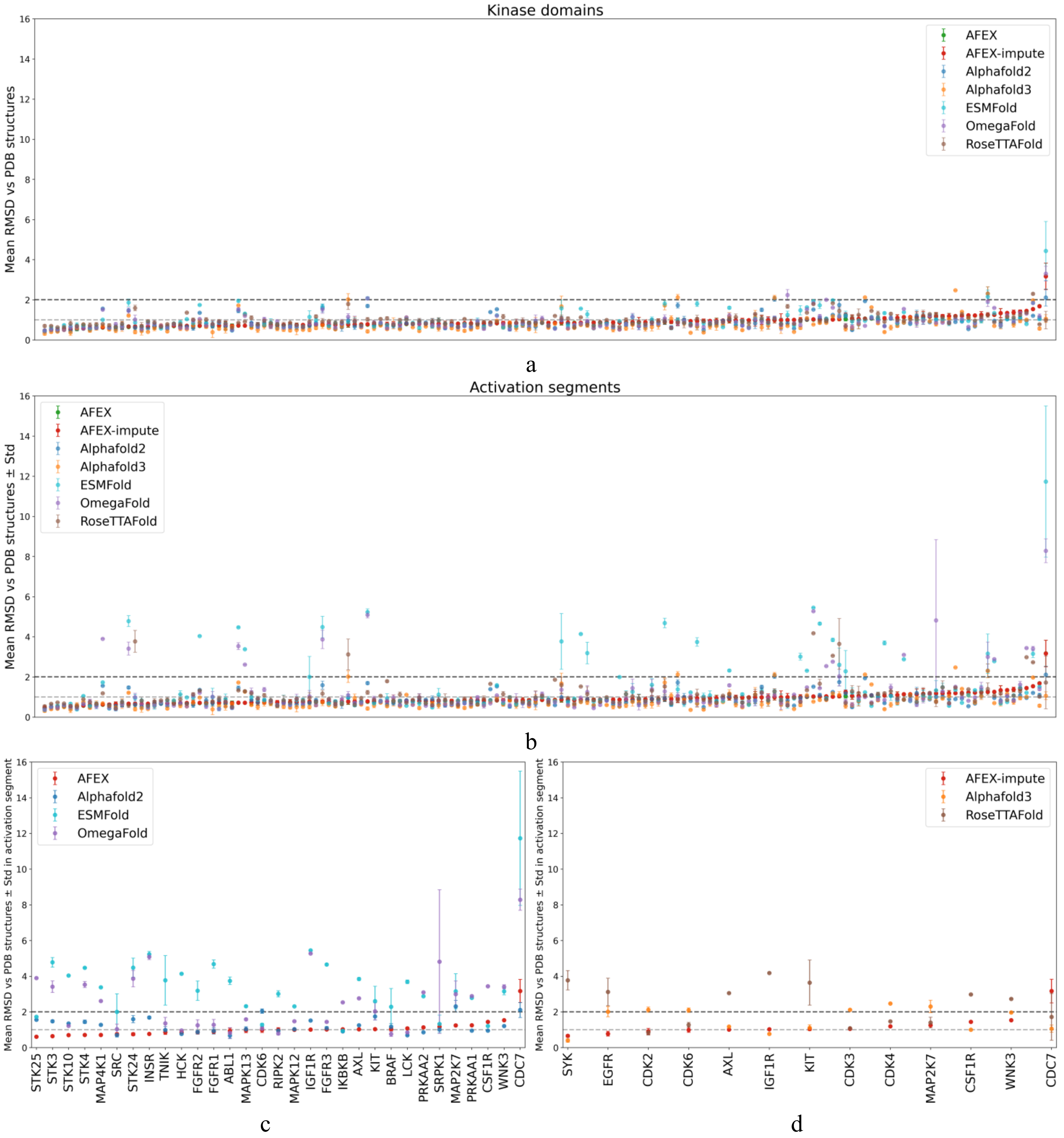


**Supplementary Figure 12. Comparison of catalytic kinase domain and activation segment prediction models with experimentally determined active, substrate-bound conformations of 156 kinases from the PDB.** Panel (a) displays the mean RMSD and standard deviation (for kinases with multiple active structures in the PDB), evaluated across all available PDB structures; results for different models are indicated by distinct colors. Panels (b)–(d) show the RMSD and standard deviation (for kinases with multiple active structures in the PDB) of activation segments, considering both all RMSD values and those exceeding 2 Å, across residues spanning from the position preceding the DFG motif to the APE motif..

**Supplementary Tables**

**Supplementary Table 1.** Mean RMSDs for active conformation of kinases excluding those with AlphaFold 2 or 3 models exhibiting RMSDs > 2 Å.

| | Mean kinase domain RMSD ($N_{kinases}$) | Mean activation loop RMSD ($N_{kinases}$) |
|---|---|---|
| AFEX vs PDB (before 2019-08-28) | 0.89 Å (144) | 0.90 Å (132) |
| AF2 vs PDB (before 2019-08-28) | 0.82 Å (144) | 0.79 Å (132) |
| AFEX vs PDB (after 2019-08-28) | 0.83 Å (65) | 0.88 Å (58) |
| AF2 vs PDB (after 2019-08-28) | 0.83 Å (65) | 0.79 Å (58) |
| AFEX vs PDB (before 2021-09-30) | 0.90 Å (146) | 0.94 Å (138) |
| AF3 vs PDB (before 2021-09-30) | 0.72 Å (146) | 0.68 Å (138) |
| AFEX vs PDB (after 2021-09-30) | 0.81 Å (39) | 0.82 Å (35) |
| AF3 vs PDB (after 2021-09-30) | 0.76 Å (39) | 0.65 Å (35) |

**Supplementary Table 2.** Quantitative statistics of active and inactive human cyclin-dependent kinase (CDK) structures in the Protein Data Bank (PDB).

| Kinase name | Active-state structures | Inactive-state structures |
|---|---|---|
| All CDKs | 393 | 782 |
| CDK1 | 0 | 28 |
| CDK2 | 318 | 618 |
| CDK3 | 0 | 0 |
| CDK4 | 1 | 16 |
| CDK5 | 18 | 1 |
| CDK6 | 5 | 16 |
| CDK7 | 4 | 12 |
| CDK8 | 0 | 48 |
| CDK9 | 28 | 11 |
| CDK12 | 16 | 31 |
| CDK13 | 3 | 0 |
| CDK16 | 0 | 4 |
